# Evolutionary transitions to self-fertilization influence the inference of introgression history

**DOI:** 10.64898/2026.05.21.726369

**Authors:** Lukas Metzger, Neda Rahnamae, Juliette de Meaux, Aurélien Tellier

**Affiliations:** Population Genetics, Department of Life Science Systems, School of Life Sciences, Technical University of Munich, 85354 Freising, Germany; Institute for Plant Sciences, Biocenter, Cologne, 50674 Germany; Institute of Plant Ecology and Evolution, Heinrich Heine University Düsseldorf, Düsseldorf 40225, Germany

## Abstract

The availability of polymorphism data and statistical inference methods allows documenting the widespread occurrence of introgression and hybridization across the tree of life. However, these methods are primarily optimized for outcrossing species without generation overlap or seed banking, thereby ignoring the consequences of life-history traits (and their evolution) on genome-wide polymorphism patterns. We investigate how a transition from outcrossing to selfing, a common feature of plant species, may affect the inference of introgression history. We simulate six demographic models with different histories of gene flow under two mating-system scenarios: a constant high selfing rate and a transition-to-selfing scenario. Using an Approximate Bayesian Computation framework with random forests, we compare model choice based on genotypic summary statistics alone and in combination with coalescent statistics derived from coalescent tree sequences. Including coalescent information substantially improves model classification, especially for distinguishing secondary contact and continuous gene flow. Cross-classification of pseudo-observed datasets shows that ignoring a transition to selfing can lead to false demographic inferences, with transition-to-selfing data often misclassified as ancient gene flow or secondary contact when analyzed under a constant selfing model. We then apply this inference framework to genomic data from *Arabis nemorensis* and *Arabis sagittata*, two predominantly selfing species with evidence for post-split hybridization. Our analyses reveal a likely transition to selfing roughly 470,000–890,000 years ago, and a likely continuous level of gene flow after the species split. The latter results lead us to revisit our previous scenario of gene flow due to secondary contact between species inferred under constant selfing. Changes in mating systems and, by extension, life-history traits can therefore bias inference about introgression if they are not explicitly modeled. Tree-sequence-based coalescent statistics provide useful information for inferring complex demographic histories that involve both gene flow and transitions to selfing.

## 1 Introduction

Introgression and hybridization may be more widespread across the tree of life than previously thought (Gilbert & Maumus, 2023; Mallet et al., 2016; Taylor & Larson, 2019). The availability of genomic data has motivated the development of new statistical inference methods to detect introgression: similarity-based (e.g., Koutsovoulos et al., 2022; Policarpo et al., 2025; Yuan et al., 2023), asymmetry-based approaches (e.g., Green et al., 2010; Lopez Fang et al., 2024; Mackintosh & Setter, 2024; Martin et al., 2014) or model-based frameworks (e.g., Excoffier et al., 2021; Flouri et al., 2019; Roux et al., 2013). There has been a large number of studies applying these new methods to different microbial, animal or plant systems (e.g., Farnitano et al., 2024; Gilbert & Maumus, 2023; Marcionetti et al., 2024; Monnet et al., 2025), highlighting the prevalence of introgression and underscoring the importance of correctly incorporating gene flow in past demographic models (Hibbins & Hahn, 2021; Komarova & Lavrenchenko, 2022). The accurate inference of the past introgression events and their rates is especially critical when investigating speciation processes involving gene flow (Monnet et al., 2025; Roux et al., 2016), as well as for estimating neutral demographic histories that serve as null models (Johri et al., 2022) for possible selection scans. However, the inference of past gene flow is impeded by possible confounding factors and other evolutionary forces. Indeed, ancient population structure (Durand et al., 2011; Mazet et al., 2016; Tournebize & Chikhi, 2024), natural selection (Smith & Hahn, 2024), or large divergence with heterogeneous evolutionary rates (Koppetsch et al., 2024; Xiong et al., 2022) do bias the inference of introgression. Although these methods often are accurate and perform well on simulated data, it becomes difficult to disentangle the contributions of these various effects in real datasets.

The theoretical understanding of how migration between populations (or species) shapes patterns of genetic variation or genealogies is still incomplete (Cousins et al., 2024; Galtier, 2024). Mackintosh and Setter (2024) described how asymmetrical gene flow influences branch lengths and how asymmetry-based methods may detect it. Generally, few theoretical expectations have been established on how the genealogies of a sample along the genome, namely the so-called ancestral recombination graphs (ARGs, (Hudson & Kaplan, 1988)), can be used to detect introgressions ((Lewanski et al., 2024; Rosenberg & Feldman, 2002), but see the recent insights in (Deng et al., 2024; Speidel et al., 2019; Whitehouse et al., 2024)). Methods based on genealogies (or coalescent trees) inference largely rely on identifying branches whose lower ends are more recent than the inferred introgression time, but whose upper ends predate the split from the introgressing group. Focusing on pairwise coalescence times enables us to detect intervals in which shared ancestry is missing between lineages within a population. As a result, two modes of coalescing events can arise and can be detected: 1) one capturing recent coalescent events in regions where all post-split ancestry remained in the recipient population, and 2) another reflecting older coalescent events in regions that are introgressed (Lewanski et al., 2024). However, it remains unclear how specific signals emerge in response to the timing, direction, and magnitude (and asymmetry) of gene flow under complex demographic scenarios involving both recent and ancient introgression.

An additional limitation is the heterogeneity in recombination and mutation rates and of the distribution of coalescent times (time to the most recent common ancestor of a sample at a locus) across the genome (Lewanski et al., 2024; Mackintosh & Setter, 2024; Rosenberg & Feldman, 2002), which may obscure and affect the reliability of introgression inference. Indeed, most methods for detecting introgression rely on simplifying assumptions about the underlying genealogy (the Kingman coalescent) and on homogeneous mutation and recombination rates across the genome, as is commonly used in methods for drawing inference of the demographic history (Deng et al., 2024; Speidel et al., 2019; Whitehouse et al., 2024). A classic assumption is that the ratio of the per population recombination (*ρ* = 4*N_e_r*; *N_e_* is the effective population size and *r* the per site mutation rate) and mutation (*θ* = 4*Nµ*; *N_e_* is the effective population size and *µ* the per site recombination rate) rates is approximately one (*ρ/θ ≈* 1 as found in humans) and has not changed in time (Strütt et al., 2023) or in the space of the genome (Sellinger et al., 2024). This assumption can be violated for many species that exhibit peculiar life-history traits that yield a ratio of *ρ/θ ̸*= 1 (Sellinger et al., 2020), and even more so for species that underwent transitions in life-history traits with concomitant temporal variation in this ratio (Metzger et al., 2026; Strütt et al., 2023).

One of the most researched and common changes in life-history trait is the transition to self-fertilization (selfing, (Stebbins, 1957)). Selfing is a life-history trait and a form of sexual reproduction in which both gametes come from the same parent. About half of all flowering plants prevent self-pollen from fertilizing the plant by a mechanism called genetic self-incompatibility (SI) (Charlesworth, 2009; Franklin-Tong, 2008). Its loss leads to increased selfing rates and has occurred many times across different plant lineages (Igic et al., 2008). This transition from outcrossing to selfing can be favored when the benefits of selfing outweigh the costs of inbreeding depression and reduced outcrossing opportunities (Abu Awad & Billiard, 2017; Abu Awad & Roze, 2020; Busch & Delph, 2011; Charlesworth, 2006; Cutter, 2019; Goodwillie et al., 2005; Igic et al., 2008). Plant species that self-fertilize at high rates often show traits distinct from their strictly outcrossing relatives. The “selfing syndrome” includes smaller flowers with closely positioned stigmas and anthers, limited pollen, and shorter flower longevity (Ornduff, 1969; Sicard & Lenhard, 2011). Selfing has a double-edged sword effect. On the one hand, the “genomic selfing syndrome” is characterized by the accumulation of harmful mutations, smaller genomes with fewer transposable elements, rapid structural changes in chromosomes, inbreeding depression, reduced diversity and stronger linkage disequilibrium (Abu Awad & Roze, 2020; Barrett et al., 2014; Glémin et al., 2019). On the other hand, selfing promotes purging of deleterious mutations, as harmful alleles become more frequently exposed (Abu Awad & Billiard, 2017; Arunkumar et al., 2014). Therefore, several key neutral and selective evolutionary processes occur during the transition from outcrossing to selfing, which are critical for the population’s fitness and survival (Abu Awad & Billiard, 2017).

When a species shifts from outcrossing to selfing, both the effective population size (*N_e_*) and the population recombination rate (*ρ*) are rescaled, which alters the genomewide pattern of variation (Nordborg, 2000; Nordborg & Donnelly, 1997). In outcrossing populations, the ratio of *ρ* to *θ* reflects *r/µ*, but selfing lowers heterozygosity and thus reduces the effective population size, and thereby also rescales (decreases) *θ*. Selfing also weakens the effect of recombination by further reducing heterozygosity, which can make crossovers less likely to create new combinations (Nordborg, 2000; Nordborg & Donnelly, 1997) (see methods for further details). Strütt et al. (2023) proposed identifying non-recombining segments that share the same most recent common ancestor (TMRCA segments). Their lengths reflect the effective population recombination rate, and the joint distribution of TMRCA and segment length can be used to detect transition to selfing. During the outcrossing phase, TMRCA segments shorten more quickly than in the selfing phase, causing a change in segment-length covariance when selfing starts. A large ancestral population alone does not explain this pattern because it also increases *θ*, producing a different TMRCA/segment-length distribution. Under a transition to selfing, segments that coalesced before the transition do appear as clustered together within chromosomes, often neighboring similarly old segments (Strütt et al., 2023). This heterogeneity in TMRCA clustering across the genome (segment length and density) is not observed under a constant life-history trait. These genomic signatures led to the development of new statistical methods (teSMC and tsABC) to estimate the age of such transitions to self-fertilization (Strütt et al. (2023), see review in Metzger et al. (2026)).

As a result, we lack a comprehensive understanding of how introgression and a transition to selfing, both of which can introduce older coalescent branches (older TMRCA) into a sample, would interact, and if and how, their footprints can be distinguished from one another. Indeed, introgressed DNA fragments may mimic the signals of transition to selfing, as both processes alter TMRCA patterns in ways that can generate shorter but older coalescence segments (Lewanski et al., 2024; Mackintosh & Setter, 2024). This can lead to elevated variation or an excess of high-frequency derived alleles, coupled with extended linkage disequilibrium. We hypothesize that when a transition to selfing is not accounted for, these characteristic regions might be misattributed to introgression, leading to misunderstandings of its timing and mode. Without models that incorporate both introgression and selfing, demographic inferences in plants may mistakenly interpret introgressed segments as evidence for a transition to selfing, causing incorrect estimates of when and how introgression occurred. To tackle these issues, we aim to (1) show how tree-sequences can help infer demographic events under a more complex speciation model, and (2) investigate how a recent transition to selfing biases introgression inference in a system where both species shifted to selfing in the past. We develop an ABC framework that tests multiple demographic scenarios and theoretically demonstrates how selfing can obscure signals of introgression. We then apply this approach to two sister plant species (*Arabis sagittata* and *Arabis nemorensis*) undergoing ongoing hybridization with secondary contact (Dittberner et al., 2022; Rahnamae et al., 2026). We first reveal that both species underwent a transition to selfing. Second, we demonstrate that accounting for the transition to selfing leads us to re-evaluate the demographic history of introgression and post-speciation contact and gene flow between these species.

## 2 Methods

### 2.1 Modelling selfing in population genomics

We assume a population with *N* diploid individuals, and selfing or outcrossing occur with probabilities *σ* and (1 *− σ*), respectively. Two main population parameters shape the genealogies along the genome, that is, their TMRCA and the length of the fragment the genealogy covers (segment of Identity by Descent) (Hudson & Kaplan, 1988; Strütt et al., 2023; Wiuf & Hein, 1999): the population mutation rate (*θ*) and the population recombination rate (*ρ*). Both depend on the effective population size and the mutation (*µ*) or recombination (*r*) rate per site per generation. In an outcrossing population, the ratio of *ρ* over *θ* equals the ratio of *r* to *µ*. However, selfing lowers heterozygosity and reduces the effective population size to *N_σ_*. We use inbreeding factor *F_IS_* based on previous theory (Fu, 1997; Golding & Strobeck, 1980; Nordborg, 2000; Nordborg & Donnelly, 1997; Padhukasahasram et al., 2008; Strütt et al., 2023), yielding:

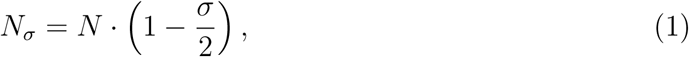

or when expressed with *F_IS_*,

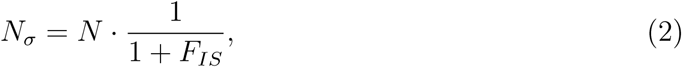

with

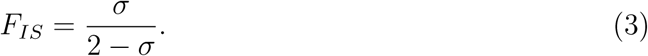

Selfing thereby scales both *θ* and *ρ* by the reduced effective population size. At the same time, selfing weakens further the effect of recombination by reducing heterozygosity, which can make crossovers less likely to create new combinations. If selfing rate is 1 (fully selfing and ignoring mutation), heterozygosity vanishes, so recombination does not change haplotypes. Thus, the recombination rate *r* is rescaled as

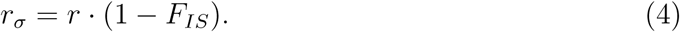

Hence, the population recombination rate *ρ* is adjusted in two ways:

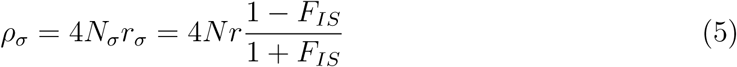

With *µ* being the mutation rate in the genome (e.g., per site), the population mutation parameter *θ* scales for the effect of selfing only by the effective population size:

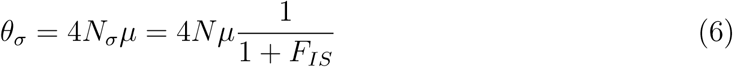

This creates a shift in the ratio

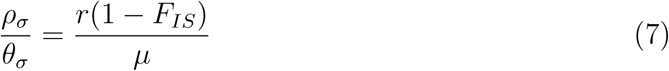

The new ratio between *θ_σ_* and *ρ_σ_* can help estimate selfing and distinguish it from outcrossing signals (Nordborg, 2000; Strütt et al., 2023). Any summary of polymorphism data that informs about both mutation and recombination can be used to infer changes in the selfing rate over time (Strütt et al., 2023).

We use a forward-in-time Wright-Fisher model (Fisher, 1930; Wright, 1931), which simulates reproduction (including recombination) from one generation to the next. This approach explicitly tracks each individual, making it well-suited for incorporating complex traits, such as selfing, and non-neutral processes. We build here on the implementation of forward-in-time simulations within the SLiM framework (Haller & Messer, 2023; Haller et al., 2019) to simulate transition to selfing.

### 2.2 Simulations and summary statistic calculation

To explore how evolutionary transitions to selfing influence the inference of introgression, we defined six demographic models based on a two-species split framework. In this setup, an ancestral population splits into two species, each of which subsequently diverges into subspecies at specific times, following a structure similar to Dittberner et al. (2022). The models included: (1) a no gene flow (GF) model; (2) an early ancient GF model where gene flow stops 100,000 generations after the species split; (3) a continuous ancient GF model with gene flow stopping after subspecies divergence; (4) a recent GF model with gene flow occurring only in the most recent times; (5) a secondary contact model with both ancient and recent gene flow separated by a period of isolation; and (6) a continuous GF model with ongoing gene flow throughout the entire timeframe.

Simulations were performed using the SLiM4 simulator (Haller & Messer, 2023; Haller et al., 2019; Kelleher et al., 2018), modeling populations under a neutral forward-in-time Wright-Fisher model. We simulated populations of diploid individuals with chromosomes of 2 Mb. Chromosome-specific recombination rates were kept constant across the 2 Mb region, modeled after chromosome 1 of *Arabis nemorensis*, with a recombination rate of 5.56 × 10^−8^. Parameters for the timing of split events, migration rates, and population sizes were drawn from uniform prior distributions. Migration rates were adjusted to be symmetrical between populations, ensuring equal rates in both directions. For population sizes, we used a log-uniform prior distribution based on the results of Dittberner et al. (2022) that suggested population sizes tend to be low. To reduce computational burden, all parameters were rescaled by a factor of 10. A total of 10,000 simulations were run for each model. An overview of the models is provided in Figure 1 and Figure 12, and the prior parameter distributions are detailed in Table 1.

**Figure 1:**
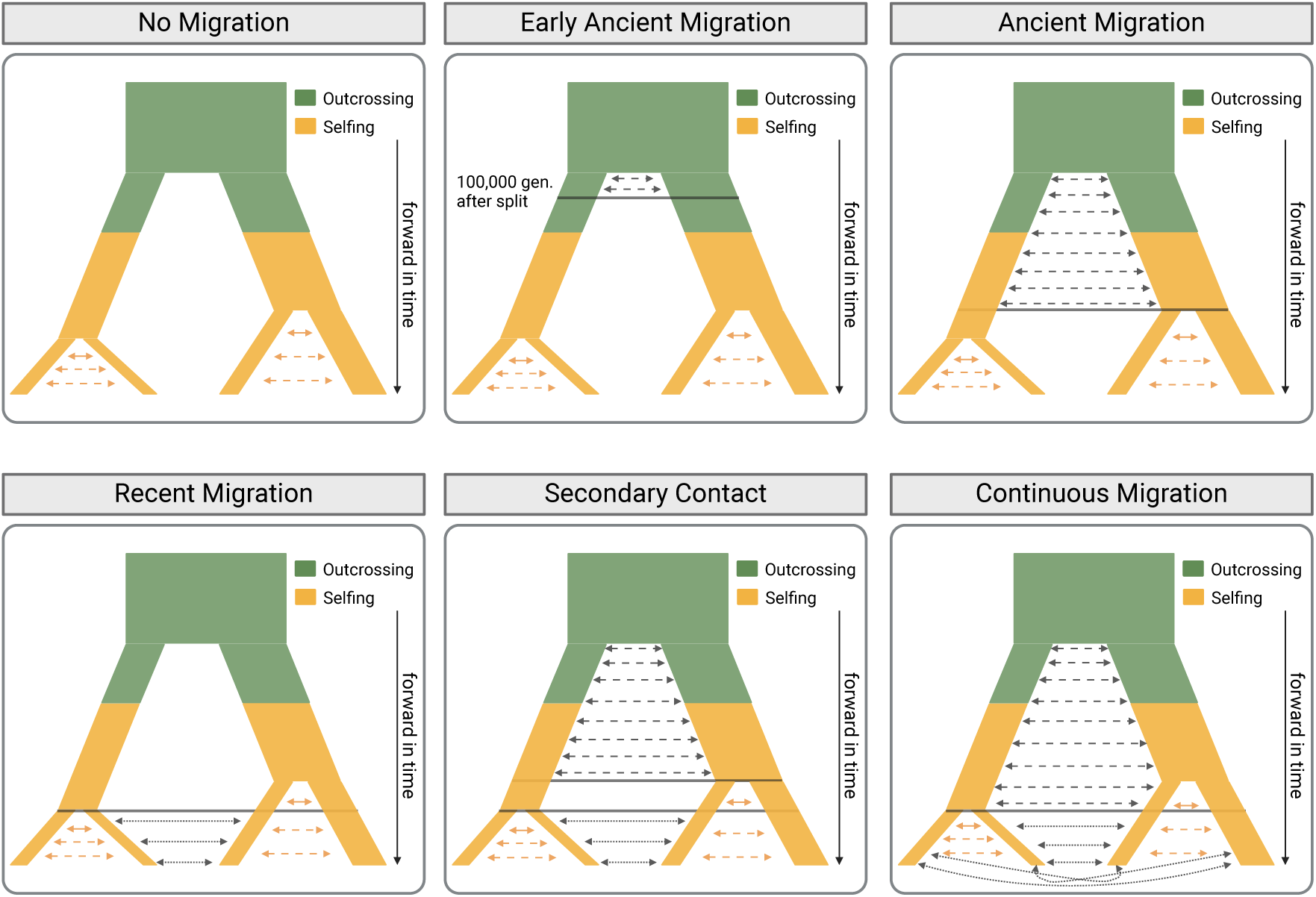
Summary of the 6 demographic models under the transition-to-selfing scenario with symmetrical gene flow. Colors represent mating systems: green denotes outcrossing, and yellow indicates selfing. Migration rates are shown with arrows, black for interspecific migration and yellow for intraspecific migration, both of which are symmetrical. Black horizontal lines indicate the timepoints of population splits or changes in migration rates. Population sizes are allowed to change after each speciation event.

**Table 1:**
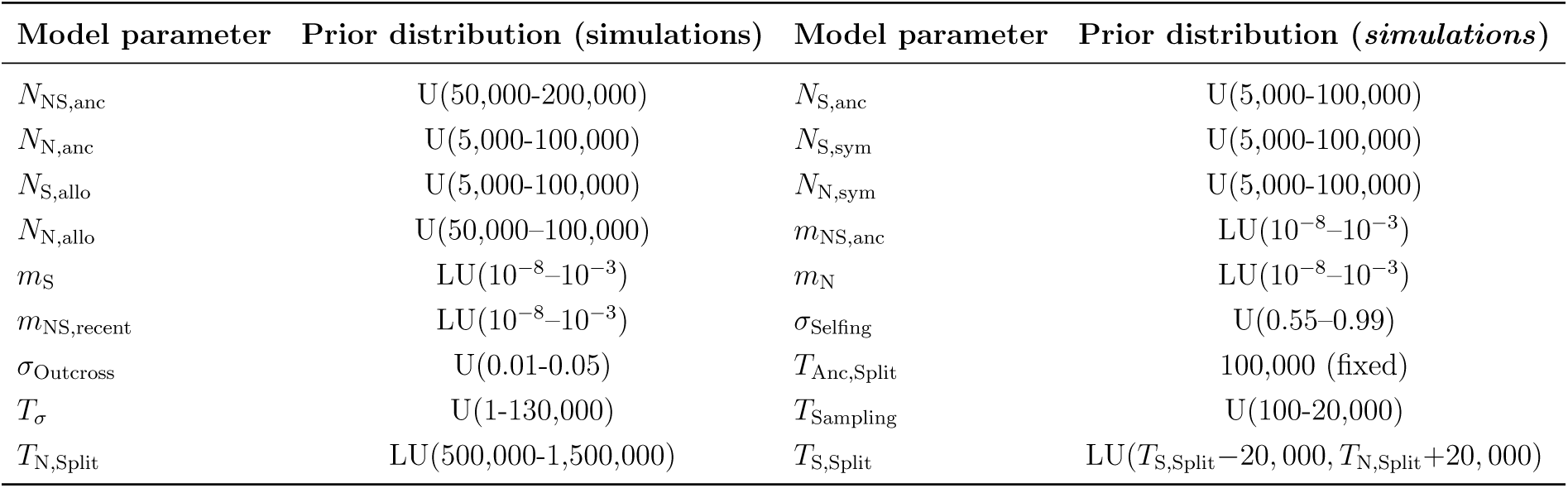
Unscaled prior distributions used to simulate data sets for the ABC. This table summarizes the unscaled prior distributions used to generate model parameters for the simulations. The priors include uniform (U) and log-uniform (LU) distributions across the parameter ranges used in the ABC. For an overview of parameter relationships, see Fig. 12. Since SLiM simulations run forward-in-time, scaling was applied to most parameters (by a factor of 10) to account for computational constraints when simulating large populations, except for migration and selfing rates, which remained unscaled. *σ*_Selfing_ and *σ*_Outcross_ were drawn from a prior distribution only for the transition scenario and kept constant at 0.9 for the constant scenario.

In one scenario, we kept the selfing rate constant at 0.9 across all six models (constant selfing scenario, CSS). In another scenario, we introduced a transition to selfing, occurring either during the ancestral population before the species split or up to 300,000 generations after the species split (*T*_AB,Split_) (transition-to-selfing scenario, TSS). Within this scenario, selfing rates were drawn from a uniform prior distribution ranging from predominantly outcrossing (*σ*_Outcross_ = 0.01–0.05) to predominantly selfing (*σ*_Outcross_ = 0.55–0.99). This transition was applied to all populations simultaneously at a specified time (*T_σ_*). We ran simulations using the tree-sequence recording option and extracted tree files (containing the full ARG) for all individuals at the end of each simulation. These files were then subsampled to retain 37 individuals, matching the real dataset of *Arabis nemorensis* and *Arabis sagittata* (Dittberner et al., 2022; Rahnamae et al., 2026).

We processed SLiM outputs with a custom Python script using the msprime, tskit, and pyslim libraries (Baumdicker et al., 2021; Kelleher et al., 2018). A neutral ”prior history” for older, uncoalesced generations (backward in time) was added using the recapitation step in pyslim. Instead of simulating forward in time until all trees coalesced, recapitation was applied to fill in the incomplete portions of the ancestral genealogy. It started at the top of the genealogies generated by SLiM and ran a coalescent simulation backward in time to complete the history of the sample’s ancestors. This ensured that genetic variation from the initial population was accurately represented. Without recapitation, the initial population would appear genetically homogeneous, underestimating the true genetic variation in the data. During this process, we rescaled the effective population size and recombination rate (as explained in Section 2.1) to account for the effects of selfing. After overlaying neutral mutations using a rescaled rate of (7 × 10^−9^) × 10 per generation per base pair, following Ossowski et al. (2010), VCF and tree files were generated for the final generation, where 37 individuals were randomly sampled from the final populations (*N_B_, sym* = 15, *N_A_, allo* = 10, *N_A_, sym* = 6, and *N_A_, allo* = 6).

We computed 65 genotypic summary statistics from each resulting VCF file and 70 coalescent summary statistics from the tree files using a custom python script resulting in a total of 135 summary statistics. The combination of both should improve the detection of signals related to transitions to selfing and introgression. The genotypic summaries included measures of genetic diversity, such as nucleotide diversity (*π*) and Watterson’s theta (*θ_w_*), as well as Tajima’s *D*. Pairwise measures like *F_ST_* and *d_XY_*were calculated to assess genetic differentiation and divergence between populations. Patterson’s *D* and *f_D_* (ABBA-BABA) statistics were determined using eight different phylogeny combinations of populations. We also calculated statistics capturing the joint site frequency spectrum (jSFS), including shared, fixed, and private polymorphisms across population pairs, as well as invariant and segregating site counts. The additional 70 coalescent summary statistics were derived from simulated tree sequence data directly or inferred with SINGER (Deng et al., 2024) for the real dataset VCF files of *Arabis nemorensis* and *Arabis sagittata*. These summaries captured mean coalescent rates, as well as the 5 % and 95 % quantiles, across three time intervals: 0–20,000 generations, 20,000–100,000 generations, and beyond 100,000 generations until complete coalescence. Additional genealogical metrics included the mean, variance, 5 % and 95 % quantiles, and skewness of the time to the most recent common ancestor (TMRCA) across populations. We also calculated statistics describing the size and variability of non-recombining regions, such as the number of trees within a segment and the mean, variance, and quantiles (5 % and 95 %) of segment size. For each 2 Mb simulation, mimicking a chromosome, we sampled 25 replicates for the 80,000 bp windows and 10 replicates for the 200,000 bp windows. This resulted in tables containing either 250,000 rows (for 80,000 bp windows) or 100,000 rows (for 200,000 bp windows). For the 200,000 bp window approach, all 100,000 rows were used directly in the ABC model choice. In the case of the 80,000 bp windows, 10 rows were randomly selected from each of the 25 replicates per simulation dataset, ensuring that sampling was stratified across replicates rather than completely random. These selected rows were then used as input for the ABC model choice analysis.

### 2.3 Whole genome re-sequencing data

Whole-genome resequencing data was used from 37 wild-sampled *Arabis nemorensis* (6 sympatric and 6 allopatric individuals) and *Arabis sagittata* (15 sympatric and 10 allopatric) individuals (Dittberner et al., 2022). Additionally, one *Arabis androsacea* individual was used as an outgroup. This data represents reviously published resequencing data for 35 *Arabis nemorensis* and *Arabis sagittata* individuals with two additional unpublished individuals that were added to this dataset. Sampling and sequencing was conducted in Dittberner et al. (2022).

Reads were mapped to a new *Arabis nemorensis* reference genome (Rahnamae et al., 2026) using BWA mem (version 0.7.17, Li and Durbin, 2009), using default parameters. Read depth and other relevant read alignment quality control metrics were computed using QualiMap v.2.2.1 (Okonechnikov et al., 2015). Average read depth across all 37 samples was 23x. Variant calling was performed using GATK version 3.8 (Poplin et al., 2018) and duplicates were marked using PicardTools (Broad Institute, 2018). Filtering was based on quality thresholds (DP *<* 20; QD *<* 2; MQ *<* 42; FS *>* 60; SOR *>* 4; ReadPosRankSum *< −*6; MQRankSum *< −*10.5). SNPs were required to be biallelic, and sites at which more than 30 % of samples were heterozygous and more than 20 % missing were discarded. *A. nemorensis* and *A. sagittata* individuals were polarized using a custom Python script with *Arabis androsacea* as an outgroup. Phasing was performed using the program shapeit2 (Delaneau et al., 2008), assuming a generation time of 1 yr and a mutation rate per generation of 7 × 10^−9^, and the recombination rates based on the genetic map produced in (Rahnamae et al., 2026). From these steps, our dataset resulted in 12,868,614 SNPs before filtering, from which we extracted 4,455,806 SNPs. When removing *Arabis androsacea* from this dataset, the VCF file contained 1,589,590 SNPs at the end.

### 2.4 Running SINGER on the *Arabis nemorensis* and *Arabis sagittata* datasets

To mimic the simulation procedure, each chromosome file was divided into 2 Mb regions, and SINGER was run separately on each region. The mutation rate (-m) was set to 7×10^−9^, the recombination-to-mutation ratio (*r/µ*, -ratio) was set to 7, and the effective population size (texttt-Ne) was set to 10,000. SINGER ran for 2000 iterations, generating 100 posterior samples with a thinning interval of 20 iterations. The inferred trees were stored as tree sequences and analyzed further using tskit (Kelleher et al., 2018). After running SINGER, we focused on 25 regions per 2 Mb region, each 80,000 bp in size, to calculate summary statistics (as explained in Chapter 2.2). SINGER’s Bayesian framework enables sampling multiple possible ARGs per window, yielding posterior samples that reflect uncertainty in ARG inference. To incorporate this uncertainty, we computed the mean of 70 coalescent summary statistics across all 100 sampled ARGs for each 80,000 bp window and used these averaged values as coalescent summary statistics.

### 2.5 Model selection using simulated data

For model selection and parameter estimation, we used an ABC approach with random forests (ABC-RF) (Pudlo et al., 2015; Raynal et al., 2018). Following Raynal et al. (2018), we performed model selection using the R package *abcrf*. We created 1,000 bootstrap samples, each containing 100,000 entries from the simulated data. For each bootstrap sample, decision trees were constructed by splitting the data based on summary statistics that best distinguished simulations from one model class compared to others. This process continued until all simulations within a bootstrap sample were assigned to a specific model class. We measured misclassification with the out-of-bag (OOB) error rate, which is the proportion of decision trees that assigned a simulation to a wrong model class, averaged over all simulations and model classes. The results were summarized in a confusion matrix, illustrating how often the predicted model matched the true model (Fig. 2). The most informative statistics that could distinguish between demographic models were stored for reference.

**Figure 2:**
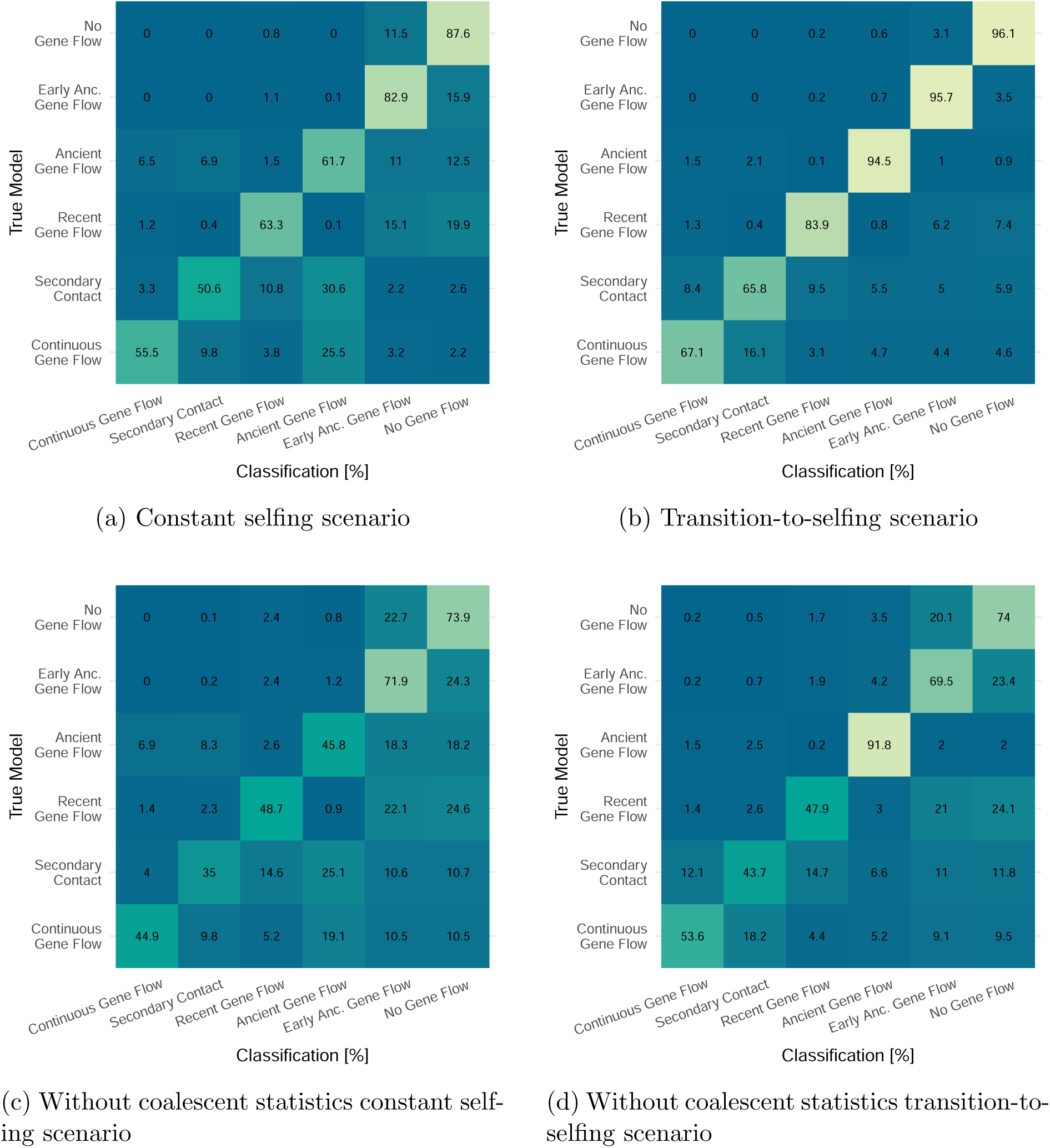
Confusion matrix for model classification using ABC-RF. The confusion matrix summarizes the classification accuracy of the ABC-RF model, comparing the predicted models (x-axis) against the true models (y-axis). Values represent the percentage of correct and misclassified instances for each model, calculated as row-wise proportions. The diagonal cells indicate the percentage of correctly classified instances for each true model, while off-diagonal cells represent misclassifications. Colors reflect classification percentages, with darker shades indicating higher accuracy. This matrix is derived from out-of-bag (OOB) predictions, which validate the model’s performance by using samples excluded from the bootstrap resampling process during model training, ensuring unbiased accuracy estimates.

To assess the effect of transitions to selfing on the accuracy of demographic model inference, we constructed pseudo-observed datasets for each demographic model and our two scenarios (TSS and CSS). For the 80,000 bp window approach, 500 randomly selected rows were sampled from each output table. This window size was chosen because from the OOB approach, the classifier performed better with narrower windows, as the 200,000 bp windows were too broad to capture all genomic signals effectively. To test the effects, we classified each pseudo-observed dataset within its own scenario and across the alternate scenario. For example, when using the classifier trained on the transition scenario (TSS), we tested its ability to correctly infer a specific pseudo-observed dataset, such as one generated under the early ancient migration model within the TSS framework. We first evaluated how accurately this dataset was classified under the transition-to-selfing scenario classifier. Then, the same pseudo-observed dataset was input into the constant scenario (CSS) classifier to assess its performance in that context. This process was repeated for each demographic model to evaluate how inference accuracy varied within and across scenarios, providing insight into potential biases introduced by transitions to selfing.

### 2.6 Model selection using the *Arabis nemorensis* and *Arabis sagittata* datasets

For the real data of *Arabis nemorensis* and *Arabis sagittata* (Dittberner et al., 2022; Rahnamae et al., 2026), we followed the same procedure as for pseudo-observed data. Summary statistics were calculated in 80,000 bp windows across all chromosomes. This dataset included 37 individuals sampled from all populations, combined into a single VCF file, which contained a total of 1,589,590 SNPs (data available from Dittberner et al., 2022; Rahnamae et al., 2026). Windows with fewer than 150 SNPs were filtered out, resulting in approximately 2,600 80,000 bp windows spanning the whole genome. The calculation of all summary statistics followed the same approach as for the simulated data, except that we used SINGER (Deng et al., 2024) to infer tree sequences and calculate coalescent statistics. SINGER, based on a Bayesian framework, allows for sampling multiple possible ARGs for a given window (we used 100 samples per window), providing a representation of the uncertainty in ARG inference. To account for this uncertainty, we calculated the mean of the 70 coalescent summary statistics across all 100 sampled ARGs for each 80,000 bp window and used these values as the coalescent summary statistics. The resulting summary statistics dataset was then used to train the classifier, which assigned the most likely demographic model to each of the 2,600 observed windows. The model most frequently selected as the best fit for these windows was deemed the most suitable for our data. We analyzed the constant and transition scenarios separately, identifying the best-fitting demographic model for each. Finally, the two best-fitting models were used to evaluate goodness-of-fit for both scenarios.

We evaluated the goodness-of-fit of the best-fitting model using a method implemented in the R package *abc* (Csillery et al., 2012), which employed a rejection algorithm. This method computed the normalized distance between the summary statistics of the observed data and all 100,000 simulated data sets. From these distances, the median of the lowest 1 % was identified during the rejection step. The same process was repeated for 1,000 simulated data sets generated under the best-fitting model, with parameter values drawn from the prior distributions (pseudo-observed data). If the model is a good fit, the observed distance (representing the 1 % best-fitting simulations for the observed data) should fall within the distribution of pseudo-observed distances (1 % best simulations fitting each pseudo-observed data set). We performed this goodness-of-fit analysis for 100 randomly chosen observed data sets. For each, a *P* -value was calculated as the proportion of pseudo-observed distances exceeding the observed distance, providing a measure of how well the model explains the data.

### 2.7 Parameter estimation in the *Arabis nemorensis* and *Arabis sagittata* datasets

For parameter estimation within a model, we used the approach described by Raynal et al. (2018). The process involved creating regression trees to group simulations with similar parameter values. This grouping was based on minimizing the squared distance (*L*^2^) between the true parameter value and the average parameter value across simulations within each group (or ”leaf”).

After identifying the best-fitting demographic model, we simulated an additional 50,000 replicates for that model under a TSS and estimated posterior distributions for each parameter. We again calculated summary statistics in 80,000 bp windows, resulting in 1,250,000 parameter sets, of which we sampled 250,000 (5 random samples within every 25 rows) to infer the parameters in the observed dataset. Parameters with log-uniform prior distributions were log-transformed to ensure uniformity, allowing all parameters to have a uniform distribution for model training.

For better prediction accuracy, we tuned the hyperparameters of the random forest models. Specifically, we tested combinations of the number of variables to split at each node (*mtry* : 11, 22, 44, 66, 88) and the minimal node size (5, 10, 30, 50). These tests were performed on all demographic parameters using the *ranger* implementation of random forests, which is also utilized by the *abcrf* package. Out-of-bag prediction errors for each combination were extracted directly from the models and summarized. Based on these results, we selected mtry = 44 and a minimal node size of 5 as the optimal settings, balancing between accuracy and training efficiency.

Using the tuned hyperparameters, we trained random forests for each demographic parameter on the full dataset of 250,000 simulations, leaving 1,000 pseudo-observed data sets aside for testing. Predictions for these test data sets were used to evaluate accuracy by comparing true and predicted values. We plotted true versus predicted values for each parameter, calculating *R*^2^ and root mean square error (RMSE). RMSE was further scaled as a percentage of each parameter’s range (%RMSE) to facilitate comparison across parameters. The theoretical 1:1 relationship between true and predicted values was visualized with a red dashed line in these plots. To assess posterior mode accuracy, we tested whether the posterior mode was more accurate than random forest estimates for certain parameters. Due to the computational intensity of posterior density estimation, this test was performed on a subset of 150 simulated datasets rather than the full 1,000 pseudo-observed datasets. We modified the *densityPlot* function to directly extract the posterior density mode, storing the true and predicted values for each parameter in a data frame. For parameter estimation from observed data, we randomly selected one of the 2600 real-data subsamples that best fit the ”continuous migration” model. Using the trained ABC-RF models, we estimated the demographic parameters for this observed data set, calculating point estimates, standard deviations, posterior modes, and the 2.5 % and 97.5 % quantiles of the posterior distributions. Log-transformed parameters were converted back to their original scale by raising them to the power of 10.

### 2.8 Estimation of selfing rate based on heterozygosity

To estimate the selfing rate based on heterozygosity estimates, we used the inbreeding coefficient (*F*) calculated from genotype data using VCFtools v0.1.17 (Danecek et al., 2011). We excluded all obvious introgressions identified in (Rahnamae et al., 2026). Specifically, the --het function was applied to the input VCF file, generating a summary for each individual, including the observed number of homozygous sites (*O*(HOM)), the expected number of homozygous sites (*E*(HOM)), the total number of genotyped sites (*N*_SITES_), and the inbreeding coefficient (*F*). The *F* values were then imported into R, where the selfing rate was calculated using the formula 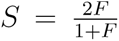, as described by Nordborg (2000). This approach allowed us to estimate each individual’s selfing rate from their heterozygosity.

### 2.9 Inference of transition to selfing using teSMC

We also used a sample of twelve haploid sequences, each from allopatric populations of *Arabis nemorensis* and *Arabis sagittata*, to estimate demography and the transition to selfing with teSMC in the one-transition mode. For this analysis, we constructed pseudo-diploid individuals by combining 12 haploid sequences from six different individuals for each species separately (6 individuals × 2 distinct haplotypes each). Regions with sequencing depth below 15, centromeric regions, and regions with introgression signals identified in Rahnamae et al., 2026 were masked. To assess the robustness of the teSMC inference across the genome, we used different genomic regions and the time-windowing option implemented in eSMC2/teSMC. These included fixed 5 Mb regions as well as additional chromosome-specific regions after masking, which varied in size and SNP number. The analyses were run with the larger predefined time-window setting in eSMC2/teSMC that uses MSMC2 time intervals (big window = TRUE). These included fixed 5 Mb regions as well as additional chromosome-specific regions after masking, which varied in size and SNP number and followed the same time-interval definition used for the MSMC2-based analyses. Different haploid sequence combinations were compared across these regions. The population size was fixed at 10,000, with recombination rates specific to each chromosome (Rahnamae et al., 2026): 5.56 × 10^−8^ for chromosome 1, 3.93 × 10^−8^ for chromosome 2, 7.79 × 10^−8^ for chromosome 3, 3.3 × 10^−8^ for chromosome 4, 5.328315 × 10^−8^ for chromosome 5, 8.16 × 10^−8^ for chromosome 6, 5.34 × 10^−8^ for chromosome 7, and 7.17 × 10^−8^ for chromosome 8. A mutation rate of 7 × 10^−9^ was used, and default parameter values were applied for all other settings.

## 3 Results

### 3.1 Model performance

To investigate how transitions to selfing affect introgression inference, we defined six demographic models based on a two-species split model (Fig. 1). In this model, an ancestral population splits into two species, each of which then diverges into populations at specific times. The scenarios are: (1) a no-migration model; (2) an early ancient migration model where gene flow stops 100,000 generations after the species split; (3) a continuous ancient migration model with gene flow stopping after the sub-population divergence; (4) a recent migration model with gene flow occurring only in the most recent times, after the last sub-population split; (5) a secondary contact model featuring both ancient and recent gene flow separated by a period of isolation; and (6) a continuous migration model with ongoing gene flow throughout the entire simulation. We drew parameters (*e.g.*, population sizes, split times, and migration rates) from prior distributions to capture a wide range of possible outcomes. Population sizes changed after each split event, and migration rates were symmetrical throughout the simulation. These models can occur under two scenarios: constant selfing (CSS) or transition-to-selfing (TSS).

As an initial evaluation, we generated 10,000 datasets for each model and each scenario, simulating a 2 Mb region using forward-in-time simulations in SLiM4 (Haller & Messer, 2023). We then applied a random-forest-based Approximate Bayesian Computation (ABC-RF) framework (Raynal et al., 2018). Using the out-of-bag (OOB) method from the abcrf package, we built confusion matrices to assess how effectively we could distinguish among the correct demographic models. In most datasets, the ABC analysis demonstrated strong discriminatory power, with high OOB classification rates. Our results indicated that including a transition to selfing improved model selection accuracy, as shown in Fig. 2a,b. Under TSS, the classifier was better at identifying the correct demographic model. In contrast, under CSS, secondary contact and continuous gene flow data were often misclassified as ancient gene flow (30 % and 25.5 % of cases, respectively). For both scenarios, the lowest classification probabilities were for the secondary contact and continuous gene flow models (50.6 % and 55.5 % for CSS, and 65.8 % and 67.1 % for TSS).

We then expanded our set of summary statistics to include coalescent statistics, not just genotypic ones. This broader approach greatly improved model selection, producing a more accurate confusion matrix. Figure 2c,d allows the comparison of ABC statistical power with the use of both genotypic and coalescent statistics to genotypic-only statistics in Fig. 2a,b). When using genotypic statistics alone, classification accuracy declined by about 10–25 % for most models. The exception was the ancient gene flow model under TSS, which could still be identified well (91.8 %) with genotypic statistics alone. Adding coalescent statistics improved its classification slightly (94.5 %). This suggests that when a transition to selfing is combined with ancient gene flow, the signal is already distinct enough to aid classification. However, using both genotypic and coalescent statistics provided a more detailed view of the data, increasing overall performance.

In ABC-RF, it is possible to extract the summary statistics that best separate demographic models. The confusion matrices from both scenarios suggest that each classifier uses different sets of statistics. Indeed, Fig. 3a,b shows that under CSS, various coalescent statistics (e.g., the 5 % quantile of the coalescent rate between the two species, and the skewness of the TMRCA distribution) rank highest. Under TSS, the top candidate is the same (the 5 % quantile of the coalescent rate between the two species), but genotypic statistics (such as theta Watterson and the number of invariant sites) also rise in importance. One reason may be that under TSS, the ancient GF model exhibits higher Watterson’s theta (Fig. 26), which likely explains why it is more accurately classified under TSS even with genotypic-only statistics. Overall, each ABC classifier uses different statistics to separate demographic models, reflecting distinct signals in the simulated data.

**Figure 3:**
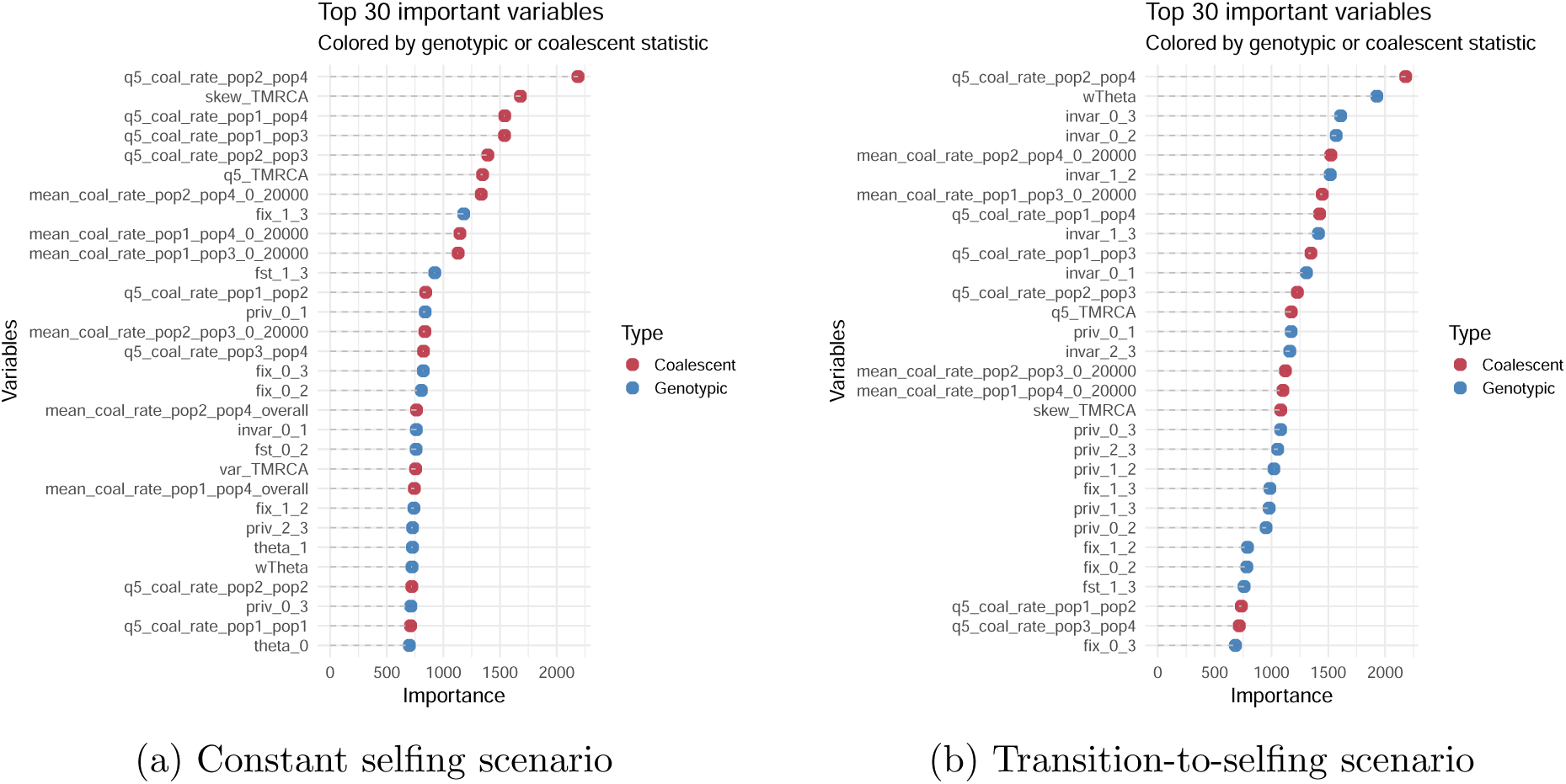
Variable importance for model classification. This plot highlights the top 30 variables that contributed most to the classification accuracy in the Approximate Bayesian Computation Random Forest (ABC-RF) framework for the no-transition set-up and the transition set-up. Variables are ranked by their importance scores, with genotypic statistics (e.g., nucleotide diversity, fixation indices, site frequency spectrum metrics) distinguished from coalescent statistics (e.g., TMRCA moments, coalescent rates across time intervals, and tree counts). The dashed lines emphasize the importance gradient for each variable, with colors distinguishing variable types.

### 3.2 Transition to selfing influences the inference of introgression

As a proof of principle, we evaluated our procedure using pseudo-observed datasets generated from an 80,000 bp window approach. This window size was chosen because in our out-of-bag (OOB) testing, the classifier performed better with narrower windows. In contrast, the 200,000 bp windows were generally too broad, resulting in slightly worse performance for most demographic models. However, for the no GF model under the CSS scenario, the 200,000 bp window captured the signals slightly better compared to the other window sizes (Fig. 17). We randomly selected 500 simulations from each demographic model, each with known input parameters, and treated their summary statistics as if they were real data. This allowed us to check whether our procedure might lead to false inferences or under- / over-estimation of key parameters under different scenarios.

To further validate our findings, we tested how the classifiers would perform when given pseudo-observed datasets from reversed scenarios (CSS vs. TSS; Fig. 4). Visually, we represented this setup by plotting each classifier (which targets one demographic model) in one figure. Each figure has two barplots (one for each classifier) and each model on the x-axis has two bars: one with red edges (representing pseudo-observed data from the opposite scenario) and one with black edges (representing pseudo-observed data from the same scenario). The bar with black edges should align with the results seen in Fig. 2a,b. When using the classifier trained on the CSS scenario to classify data simulated under the transition-to-selfing scenario, we observed erroneous assignments, particularly for the secondary contact and continuous gene flow models (Fig. 4e,f, top panel). For instance, the bars with red edges in the top panel of Fig. 4e showed that when the classifier is trained under CSS but tested on secondary contact data generated under TSS, it gave many random forest votes to both the Secondary Contact and Ancient GF models. However, most pseudo-observed parameter sets were ultimately classified as Ancient GF, even though they originated from a Secondary Contact model.

**Figure 4:**
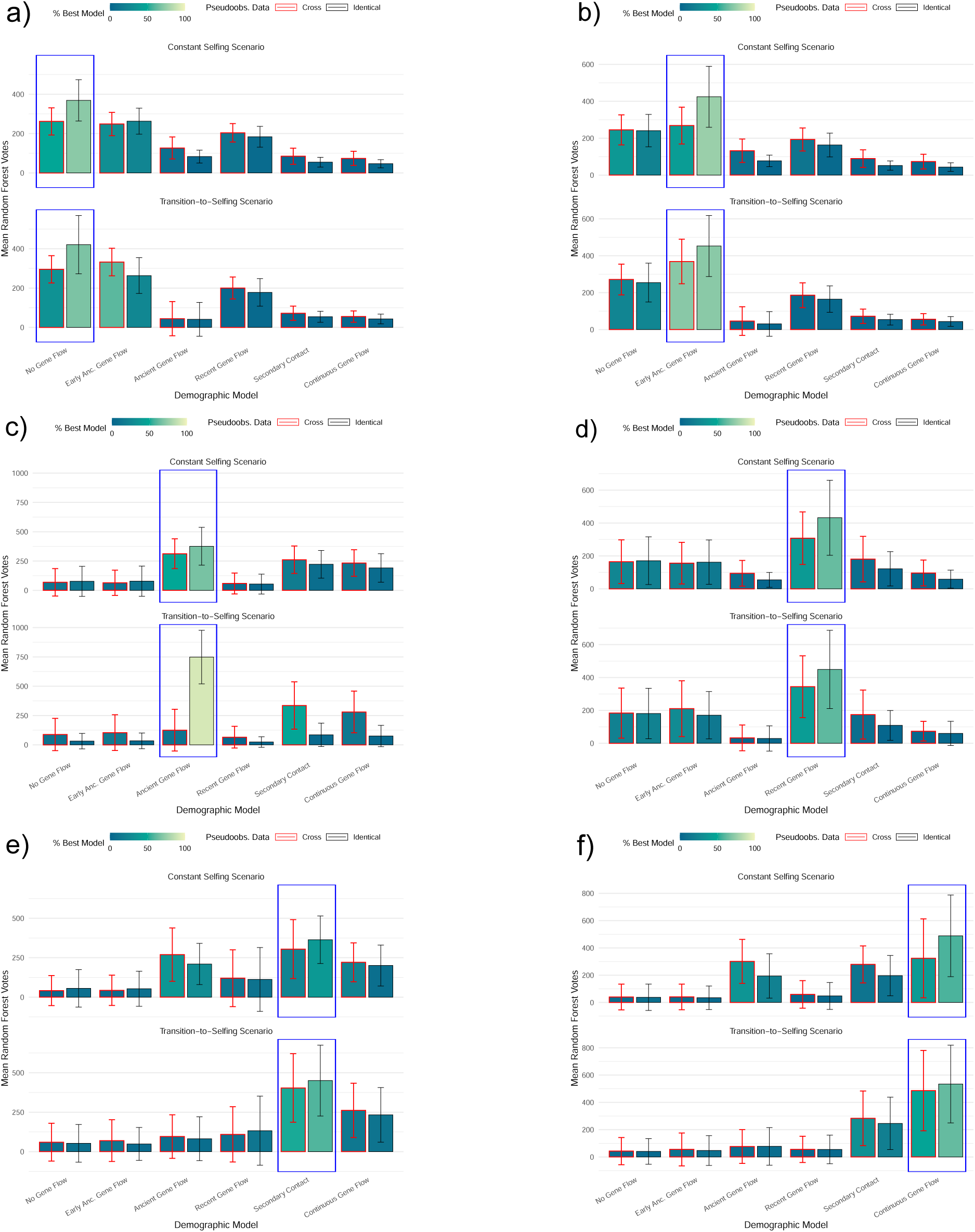
Classification performance of demographic models using pseudoobserved data under CSS and TSS. Bars represent the mean RF votes for each model, with error bars showing standard deviations. Results are categorized based on whether the pseudoobserved data originated from the same scenario (”Identical”; in black) or a different scenario (”Cross”; in red). The fill gradient indicates the percentage of best model votes. Blue box indicates the current demographic model from which the pseudoobserved data originated. Each barplot exists two times, the top barplot within each plot shows the ABC classifier trained under CSS and the bottom was trained under TSS. This plot shows the classification abilities of no GF (a), early ancient GF (b), ancient GF (c), recent GF (d), Secondary Contact (e) and continuous GF (f) models.

Conversely, the classifier trained on TSS (Fig. 4e, bottom panel, bars with red edges) more accurately disentangled the models, even when applied to CSS datasets, by correctly inferring Secondary Contact for pseudo-observed data that actually came from CSS. A similar pattern appeared in Fig. 4b,f, where the CSS-trained classifier struggled to detect the correct demographic model for pseudo-observed data derived from TSS, while the reversed setup was more accurate.

However, the opposite is true for the no gene flow (no GF) and ancient gene flow (ancient GF) models (Fig. 4a,c). In these cases, the TSS classifier failed to assign the correct model when the data stemmed from CSS. For no GF, the TSS classifier erroneously assigned pseudo-observed datasets as an early ancient GF model. For the ancient GF model, the TSS classifier often labeled it as a Secondary Contact model, with the Ancient GF model ranking only third.

This asymmetry underscored the risk of false positives and false inferences when the true mating system is not accounted for. If a transition to selfing occurred recently (and if the timing and amount of gene flow is unknown), the wrong model can be erroneously favored in ABC-based demographic inference. Neglecting such transitions can lead to misclassification and incorrect conclusions about gene flow and population history.

### 3.3 Selfing rate and transition to selfing in two *Arabis* species

We revisit previous demographic and species split analyses from (Dittberner et al., 2022; Rahnamae et al., 2026) for the two species *Arabis nemorensis* and *Arabis sagittata*, which show a history of introgression and gene flow. For *A. nemorensis* and *A. sagittata*, both species are known to be predominantly selfing (Dittberner et al., 2022), and both lack functional S-loci based on their reference genomes. We calculated selfing rates based on observed vs. expected heterozygosity within individuals, finding values ranging from 0.6 to 0.99 (see Fig. 5). When combining both allopatric populations, the estimated selfing rates ranged from 0.9 to 0.99. Indeed, when both species are pooled, the number of homozygous SNPs increases more than the number of heterozygous sites as many SNPs are fixed within each species but differ between them. Thus, this rate also reflected the number of heterozygous SNPs per population. *A. sagittata* had the highest number of heterozygous sites and thus the lowest estimated selfing rate among our samples. Note that in the allopatric *A. sagittata* population, one individual had a low selfing rate (and thus high heterozygosity) and was excluded from further analysis.

**Figure 5:**
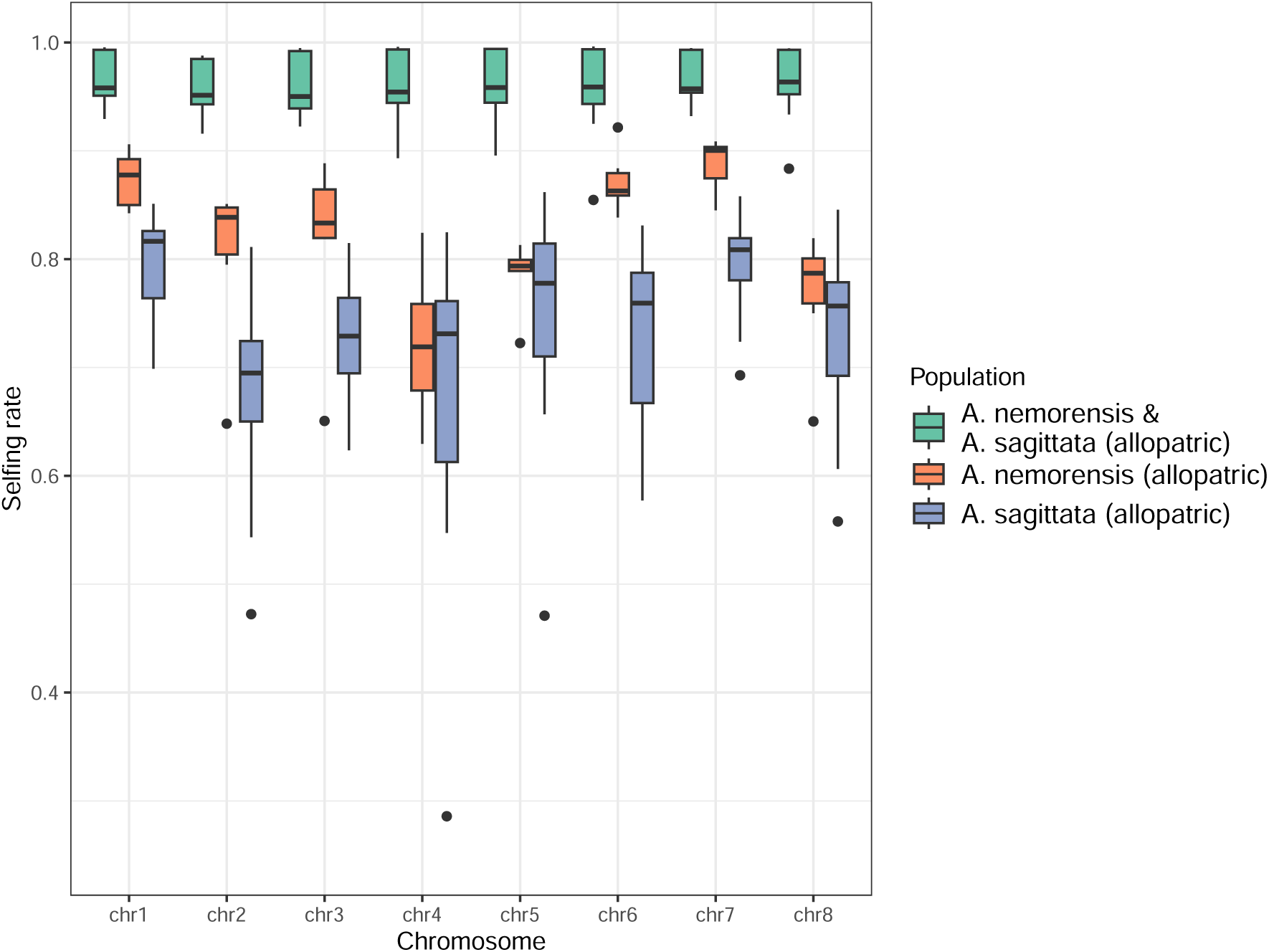
Boxplot of selfing rates across chromosomes for different populations. The populations include *A. nemorensis* and *A. sagittata* together (green), *A. nemorensis* alone (red), and *A. sagittata* alone (blue). Selfing rates were estimated based on heterozygosity for each individual, using six allopatric individuals from *A. nemorensis* and ten from *A. sagittata*. Each estimate reflects the selfing rate of a single individual, calculated based on the number of SNPs in the VCF file.

We applied teSMC to allopatric pseudo-diploid individuals to avoid biases from hybridization in sympatric populations. We selected genomic regions with coverage above 15, excluding centromeric regions and known introgressions. This yielded three regions per chromosome: one region of 5Mb, plus two other regions with sizes varying by chromosome. For each region (3 regions x 8 chromosomes), we created two pseudo-diploid combinations. We also considered different species groupings: (1) combining both species, (2) *A. nemorensis* alone, and (3) *A. sagittata* alone. The Figures 19–25 summarized all results and SNP counts. The analysis in Fig. 6 showed an effective population size of about 1,000,000 around 470,000–600,000 generations ago (one generation = one year), followed by a decline to about 10,000 approximately 1,000 years ago (younger times are less reliably inferred by SMC methods). The timing of the transition to selfing coincided with the population size decrease, around 500,000 years ago, and each chromosome showed a selfing event at that time, with selfing rates ranging from 0.6 (chromosome 2) to 0.9 (chromosomes 3 and 8). This aligned with the heterozygosity-based selfing rate estimates (0.6–0.99 in Fig. 5). Transition events occurred at different times depending on the chromosome, with the most recent estimate for chromosome 4 at approximately 470,000 years ago and the oldest estimate for chromosome 8 at around 890,000 years ago.

**Figure 6:**
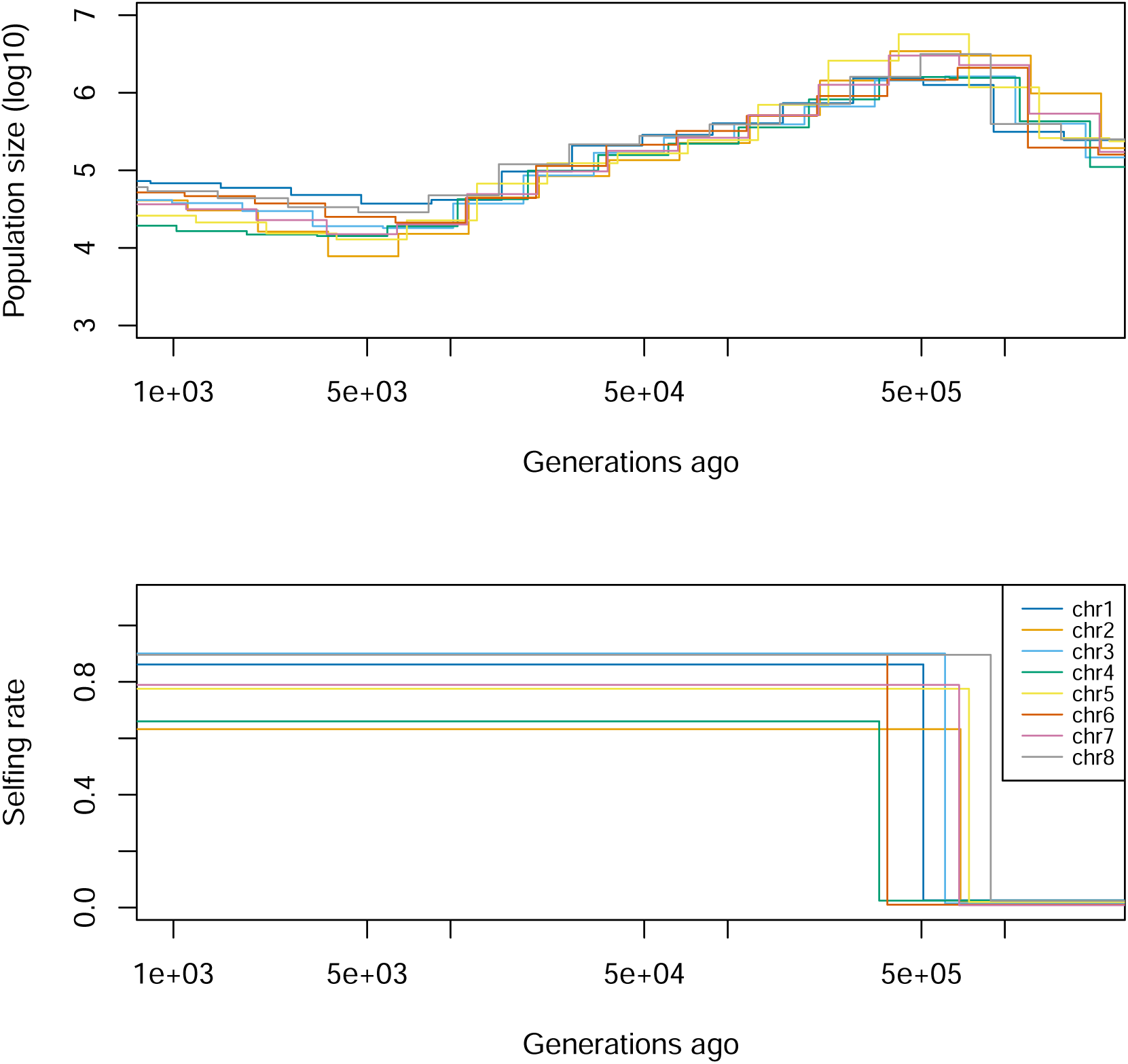
Demographic inference and transition from outcrossing to selfing in allopatric *A. nemorensis* and *A. sagittata*. The top panel illustrates estimated population sizes over time, while the bottom panel highlights transition times and selfing rates inferred for 5 Mb regions across all chromosomes in both species using teSMC. Population sizes were derived using MSMC2 time intervals within teSMC. The analysis included 12 pseudo-diploid individuals, combining two haploid sequences from distinct individuals of the same species, with 12 haploid samples from each *A. nemorensis* and *A. sagittata*. Each chromosome is represented by a colored line.

In the other datasets, when *A. nemorensis* and *A. sagittata* were analyzed separately, some regions had too few SNPs to confidently infer a transition, leading to convergence issues in teSMC (see supplementary figures Fig. 19–25).

### 3.4 Model choice results in the *Arabis* species

Using the random-forest classifier trained under TSS, we selected the best model for 2,600 observed datasets, each consisting of a single 80 kb locus. Models with non-constant gene flow consistently received the most votes. Among these, continuous gene flow was chosen as the best model for 85 % of the loci (Fig. 7), followed by secondary contact (12 %) and recent gene flow (3 %). The other three models were never identified as the most likely. These results differ from our previous analysis in Dittberner et al. (2022), where it was concluded that these species experienced an ancestral and recent sympatric migration phase separated by isolation (secondary contact model). Our new findings suggested that, when we include coalescent statistics under a transition-to-selfing scenario, gene flow appeared to have been continuous.

**Figure 7:**
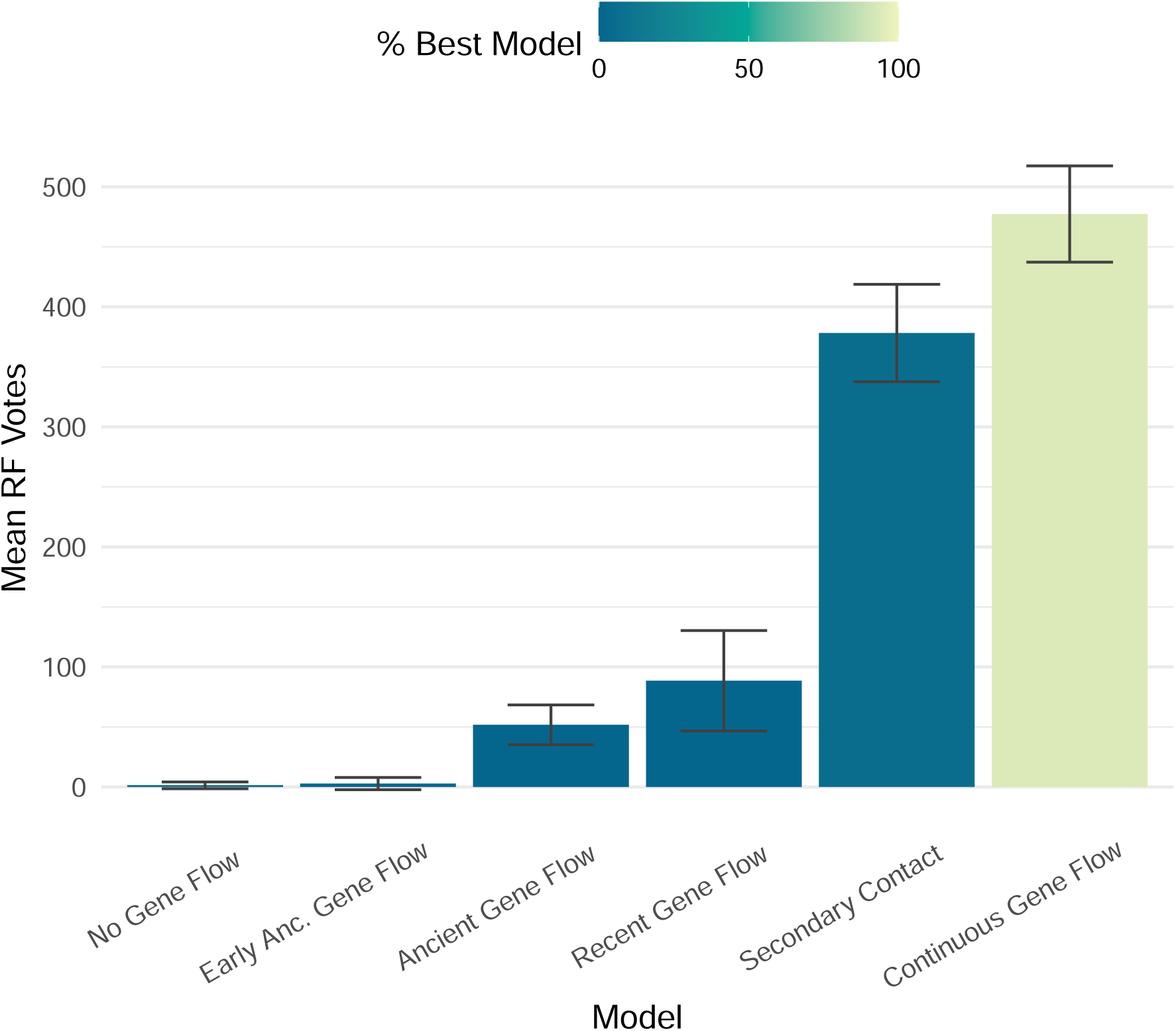
Mean random forest votes for observed data sets under the transition scenario. The bar plot displays the mean random forest votes for each demographic model, calculated from 1,000 bootstrap trees and based on 2,600 genomic windows across the observed data. Confidence intervals are shown above each bar, derived from the distribution of random forest votes across bootstrap replicates. Bars are colored to indicate the proportion of windows (%) where each model was classified as the best fit.

Applying the constant-selfing classifier to the 2,600 observed loci yielded a similar outcome: continuous gene flow remained the top choice (68%), but the secondary contact model was chosen 26 % of the time, and ancient gene flow ranked higher (4 %). Overall, the result still supported continuous gene flow as the most plausible scenario for these species.

To confirm the fit of the continuous gene flow model under TSS, we measured the distance between the observed and simulated datasets and found no significant deviation from zero, indicating that the model expectations matched rather well the observed data.

Finally, as a proof of principle, we used only genotypic statistics under constant selfing to classify the 2,600 observed datasets, similar to the approach of Dittberner et al. (2022).

In this setup, the classifier performed worse (Fig. 10). Although the continuous gene flow model was still selected in about 45 % of cases, the secondary contact model increased to 33 %, and the ancient gene flow model rose to 21 %. This spread in classifications was less robust than our earlier results under the transition scenario.

### 3.5 Parameter estimation for the demographic models for the *Arabis* species

Using the transition-to-selfing scenario with continuous migration, we next estimated the timing and intensity of gene flow, the timing of the transition event, and the sizes of ancestral and contemporary populations. We simulated 250,000 parameter sets under this demographic scenario and applied a random-forest-based ABC for parameter estimation (Raynal et al., 2018), which provides confidence intervals. Each parameter was estimated independently. To test estimation accuracy, we used 1,000 simulations excluded from the training set. The highest accuracy was achieved for the new selfing rate after the transition (*SelfingRateNew*) and the ancient migration rate between species (*M NSAnc*), whereas lower accuracy was observed for the migration rates between sympatric and allopatric populations (*mN* and *mS*; Fig. 11).

We then analyzed observed data using one randomly chosen parameter set assigned to the selected model. The estimated post-transition selfing rate was 0.81 (SD = 0.063), and the transition event occurred at 1,464,935 generations ago (SD = 88,437). This timing is about three times older than the estimate from teSMC. For *A. nemorensis*, contemporary population sizes were estimated at 20,775 (SD = 12,981) in the allopatric population and 17,044 (SD = 12,057) in the sympatric population (Fig. 12). For *A. sagittata*, these sizes were 40,412 (SD = 13,516) and 39,844 (SD = 15,172), respectively. These findings aligned with previous results showing *A. sagittata* has higher genetic diversity (Dittberner et al., 2022). The allopatric populations are larger than the sympatric ones, likely because the allopatric samples include individuals from multiple sites. Compared to Dittberner et al. (2022), our approach yielded larger population sizes, and a much older transition estimate than teSMC. One reason may be that the chosen 80 kb region has small non-recombining segments, many coalescent trees within one window, and older TMRCAs, making it more likely to reflect an earlier transition event.

We inferred the ancestral population of *A. sagittata* (79,137, SD = 11,159) to be larger than that of *A. nemorensis* (60,057, SD = 15,806). A similar decline has been seen in selfing species like *Arabidopsis thaliana* (Durvasula et al., 2017). However, *A. thaliana* has contemporary population sizes at least an order of magnitude larger, suggesting *A. nemorensis* and *A. sagittata* faced stronger population-size reductions. Our teSMC estimates of population sizes were also within a similar range.

The divergence between *A. nemorensis* and *A. sagittata* was estimated at 1,466,055 generations ago (SD = 50,313), with an ancestral size of 175,540 (SD = 11,738). Notably, this timing matched the estimated transition event, which could lead to confounding signals if the simple speciation model resembles a bottleneck-driven reduction in population size (Strütt et al., 2023). Each species then splitted into allopatric and sympatric populations around 17,113 generations ago (SD = 9,584) for *A. nemorensis* and 15,719 generations ago (SD = 4,606) for *A. sagittata*, aligning with the last glacial period. These estimates suggested that both species formed distinct subpopulations after the speciation event.

The analysis revealed low but higher migration rates than in Dittberner et al. (2022), supporting continuous gene flow as a better fit than secondary contact (Fig. 7). Migration rates were reported in log_10_-transformed proportions of individuals migrating per generation. Because we modeled symmetrical gene flow (both directions simultaneously), the results likely overestimated actual migration. We found the ancestral gene flow rate (−6.303, SD = 0.53) was greater than the recent rate (−5.50, SD = 0.98). Both were higher than in Dittberner et al. (2022), possibly due to symmetrical migration. Other migration estimates were similar to the recent interspecific rate (−5.49, SD = 1.58 for *A. sagittata* and −5.81, SD = 1.29 for *A. nemorensis*).

### 3.6 Identifying footprints of transition to selfing in *Arabis nemorensis* and *Arabis sagittata*

We next combined both the constant-selfing and transition-to-selfing scenarios under continuous gene flow in a single ABC framework to assess which model better explained each of our 2,600 parameter sets. The results in Fig. 7 and Fig. 8 indicated that both models favored continuous gene flow, so comparing them directly would allow us to pinpoint regions that may show signals of a transition.

**Figure 8:**
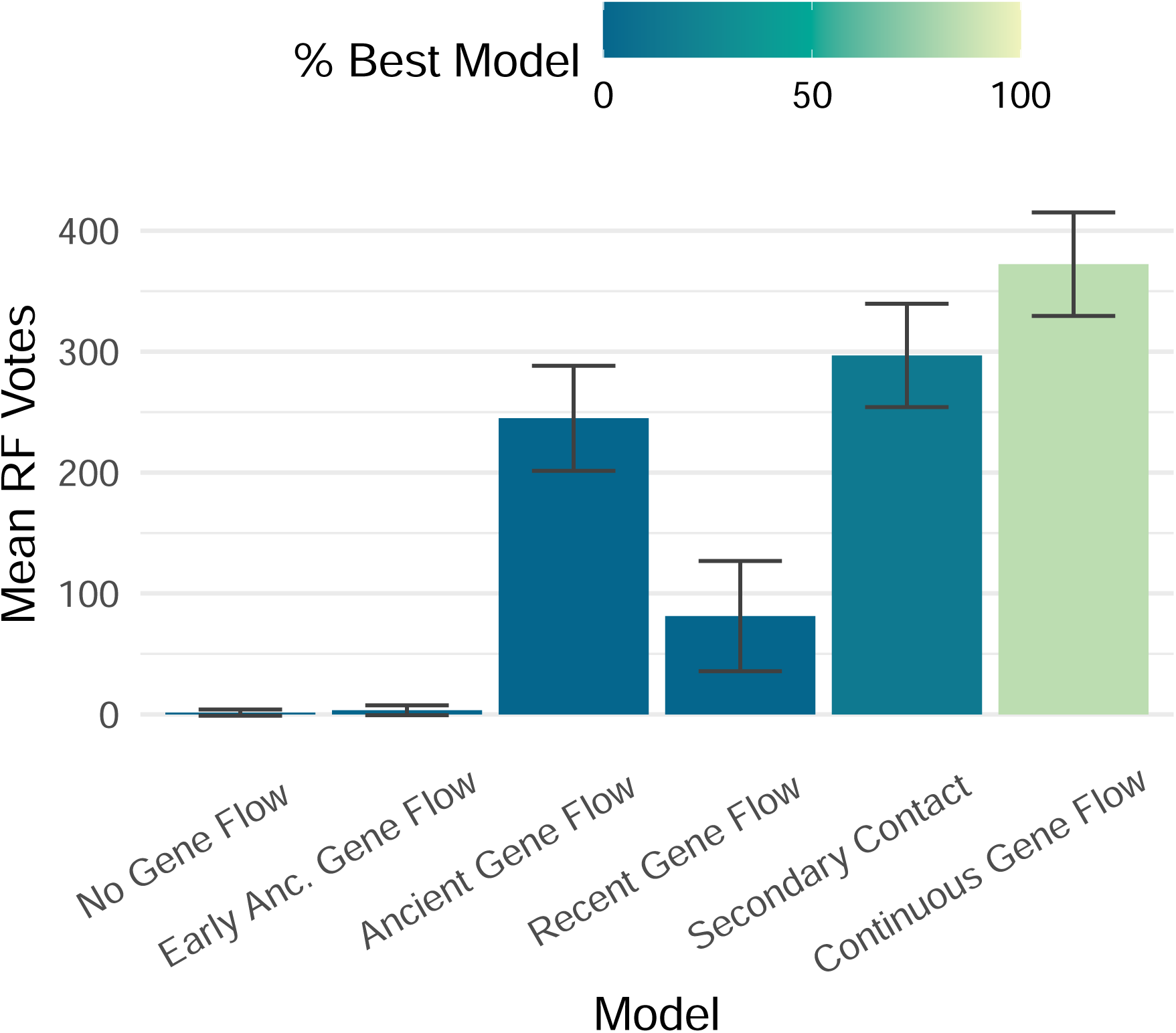
Mean random forest votes for observed data sets under the constant selfing scenario. The bar plot displays the mean random forest votes for each demographic model, calculated from 1,000 bootstrap trees and based on 2,600 genomic windows across the observed data. Confidence intervals are shown above each bar, derived from the distribution of random forest votes across bootstrap replicates. Bars are colored to indicate the proportion of windows (%) where each model was classified as the best fit.

**Figure 9:**
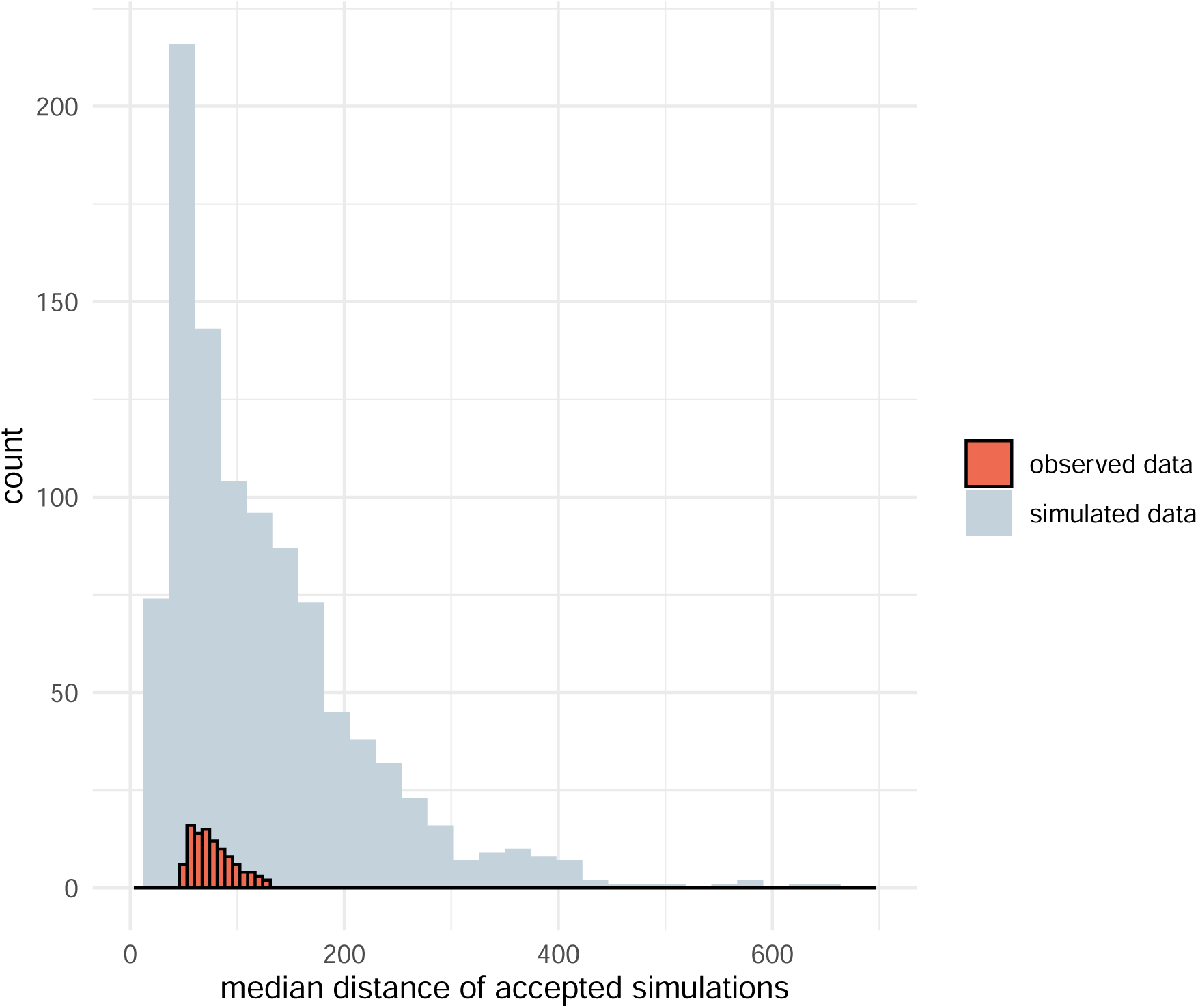
Goodness of fit for the observed data. The histogram shows the distribution of median normalized distances for 1,000 pseudo-observed data sets generated under the best-fitting model. These distances represent the closest 1 % of simulated data sets to each pseudo-observed data set (grey bars). The red bars indicate the corresponding median distance for the 100 observed data sets, providing a comparison to assess how well the model fits the observed data.

**Figure 10:**
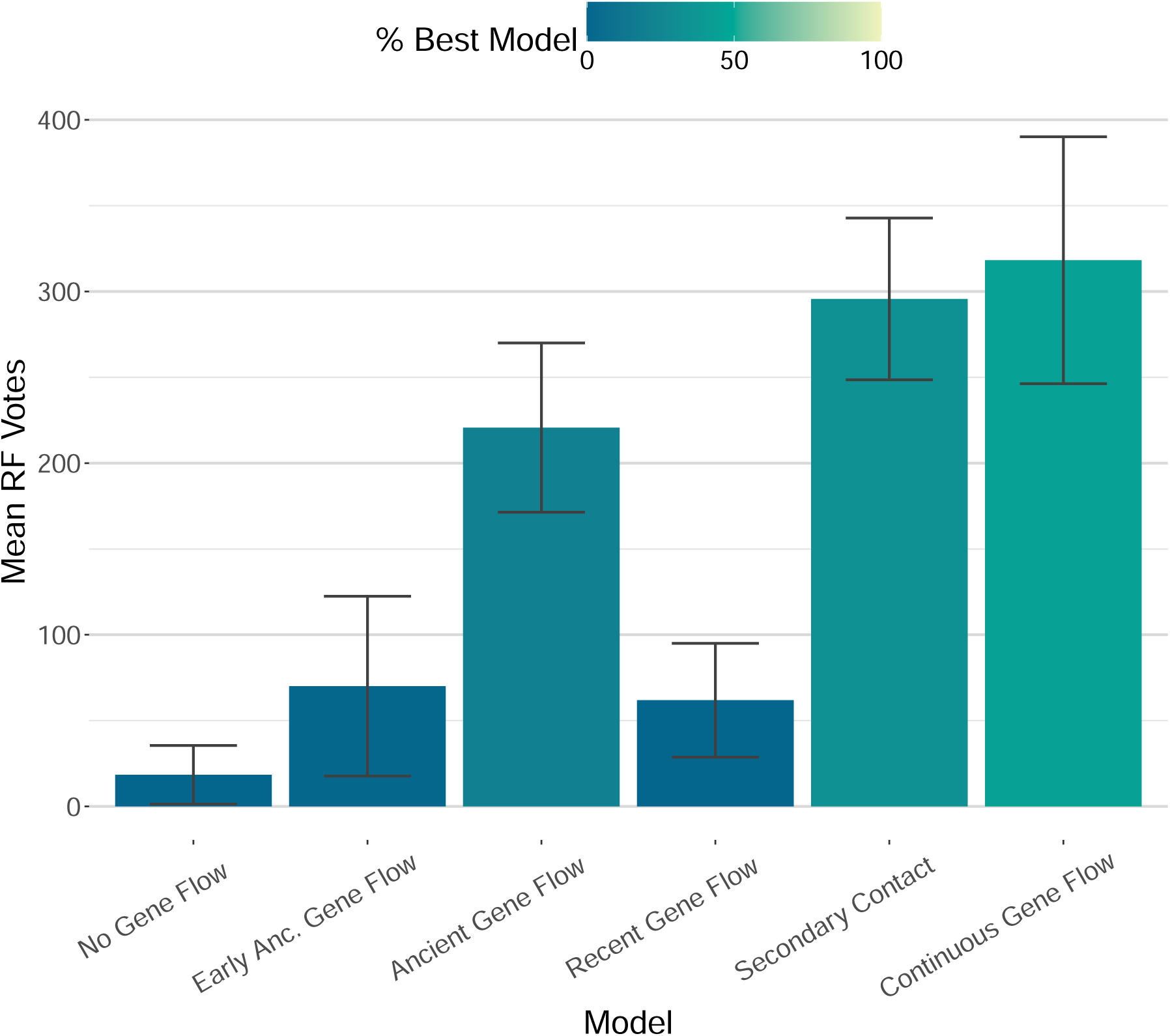
Mean random forest votes for observed data sets under the constant scenario using genotypic statistics. The bar plot displays the mean random forest votes for each demographic model that was simulated under the constant selfing scenario as in Dittberner et al. (2022), calculated from 1,000 bootstrap trees and based on 2,600 genomic windows across the observed data. Confidence intervals are shown above each bar, derived from the distribution of random forest votes across bootstrap replicates. Bars are colored to indicate the proportion of windows (%) where each model was classified as the best fit.

**Figure 11:**
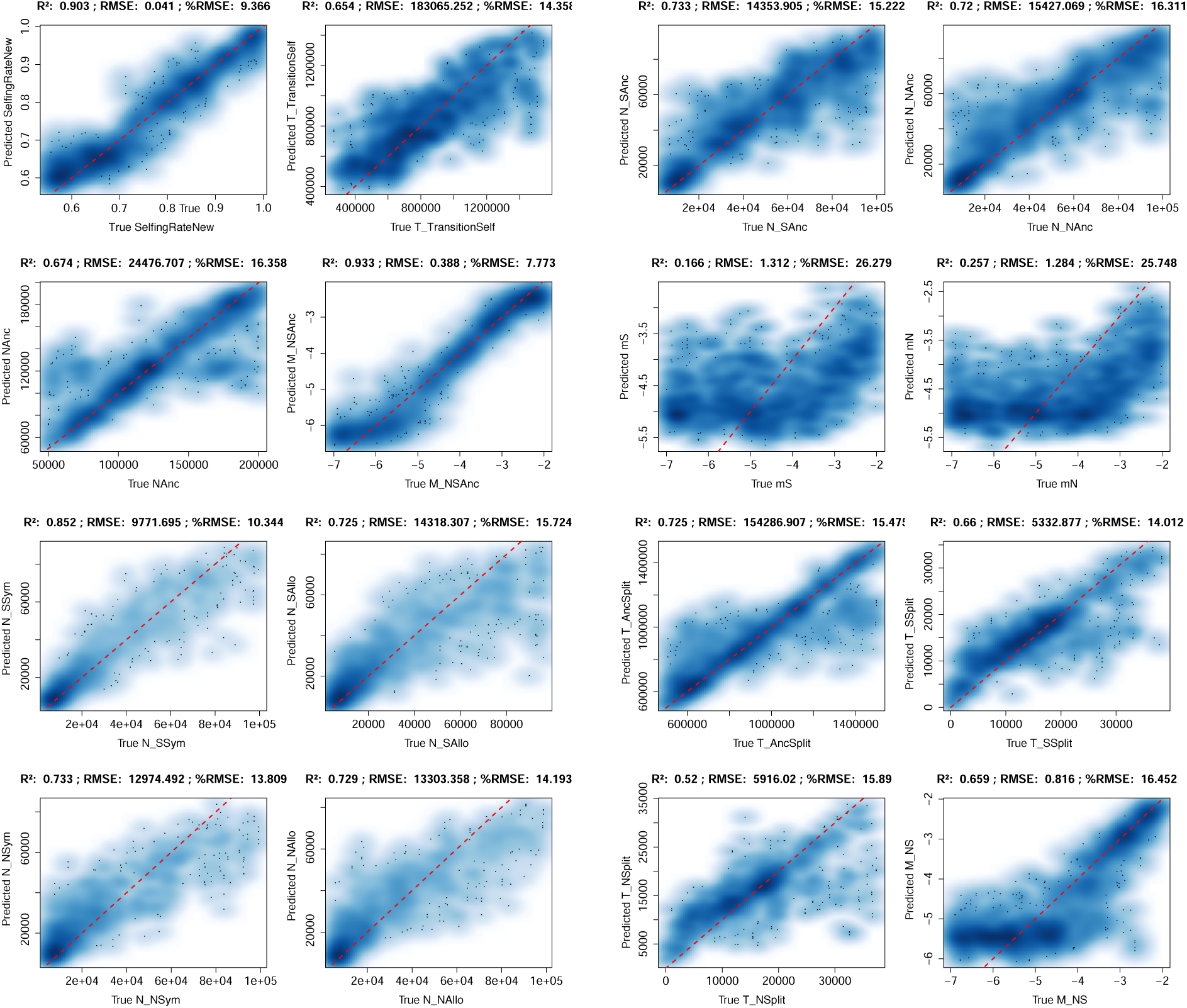
Comparison of true and predicted demographic parameters across simulations. To evaluate prediction accuracy, demographic parameters were estimated for 1000 simulations not included in the training data. True versus predicted values are shown for each parameter, along with *R*^2^ and the root mean squared error (scaled as a percentage of the parameter range). The red dashed line indicates the theoretical 1:1 relationship. Parameter names differ from those in the main text: *SelfingRateNew* – Selfing rate after the transition; *T _TransitionSelf* – Time of the transition event; *NAnc* – Ancestral population size prior to speciation; *M _NSAnc* – Migration rate between the ancient populations of *A. nemorensis* and *A. sagittata*; *N _SAnc* –Effective population size of the ancient *A. sagittata*; *N _NAnc* – Effective population size of the ancient *A. nemorensis*; *mS* – Migration rate between sympatric and allopatric *A. sagittata* populations; *mN* – Migration rate between sympatric and allopatric *Arabis nemorensis* populations; *N _SSym* – Effective population size of sympatric *A. sagittata* population; *N _SAllo* – Effective population size of allopatric *A. sagittata* population; *N _NSym* – Effective population size of sympatric *A. nemorensis* population; *N _NAllo* – Effective population size of allopatric *A. nemorensis* population; *T _AncSplit* – Time of speciation; *T _SSplit* – Time since split of ancient *A. sagittata*; *T _NSplit* – Time since split of ancient *A. nemorensis*; *m_NS* – Recent migration rate between species.

**Figure 12:**
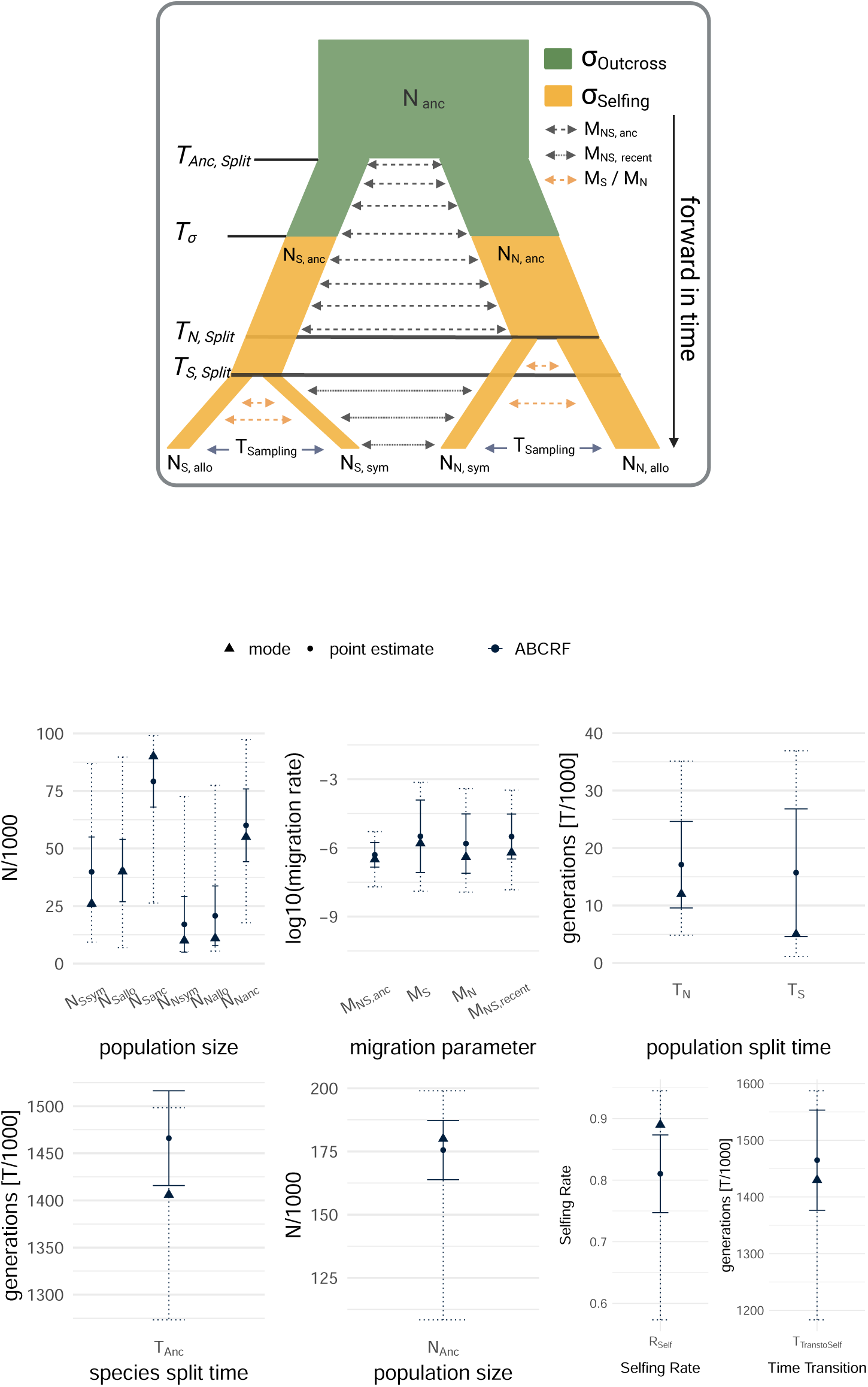
Model parameter estimation. (Top panel) Diagram illustrating the model parameters (*N* : effective population size; *M* : migration rate, proportion of migrants per generation; *T* : species/population split times or transition time) and their estimation using all summary statistics. (Bottom panel) Parameter estimates for *Arabis nemorensis* and *Arabis sagittata*, derived from the ABC-RF method. Circles represent point estimates, triangles indicate the mode of ABC-RF posterior distributions, and error bars denote standard deviations (solid) and 95 % prediction intervals (dashed).

Our ABC could distinguish between these two scenarios reasonably well, with 99.6 % of simulations correctly identified for continuous gene flow under constant selfing and 96 % for continuous gene flow under transition-to-selfing (Fig. 13). The most important statistics differentiating these scenarios were all coalescent-based (Fig. 14), including the 5 % quantile of non-recombining segment size, the number of recombination breakpoints (tree count) within each window, the 95 % quantile of TMRCA, and the variance of TMRCA. This aligned with Strütt et al. (2023), who showed that only transition-aware metrics or their derivatives can reliably distinguish transition events.

**Figure 13:**
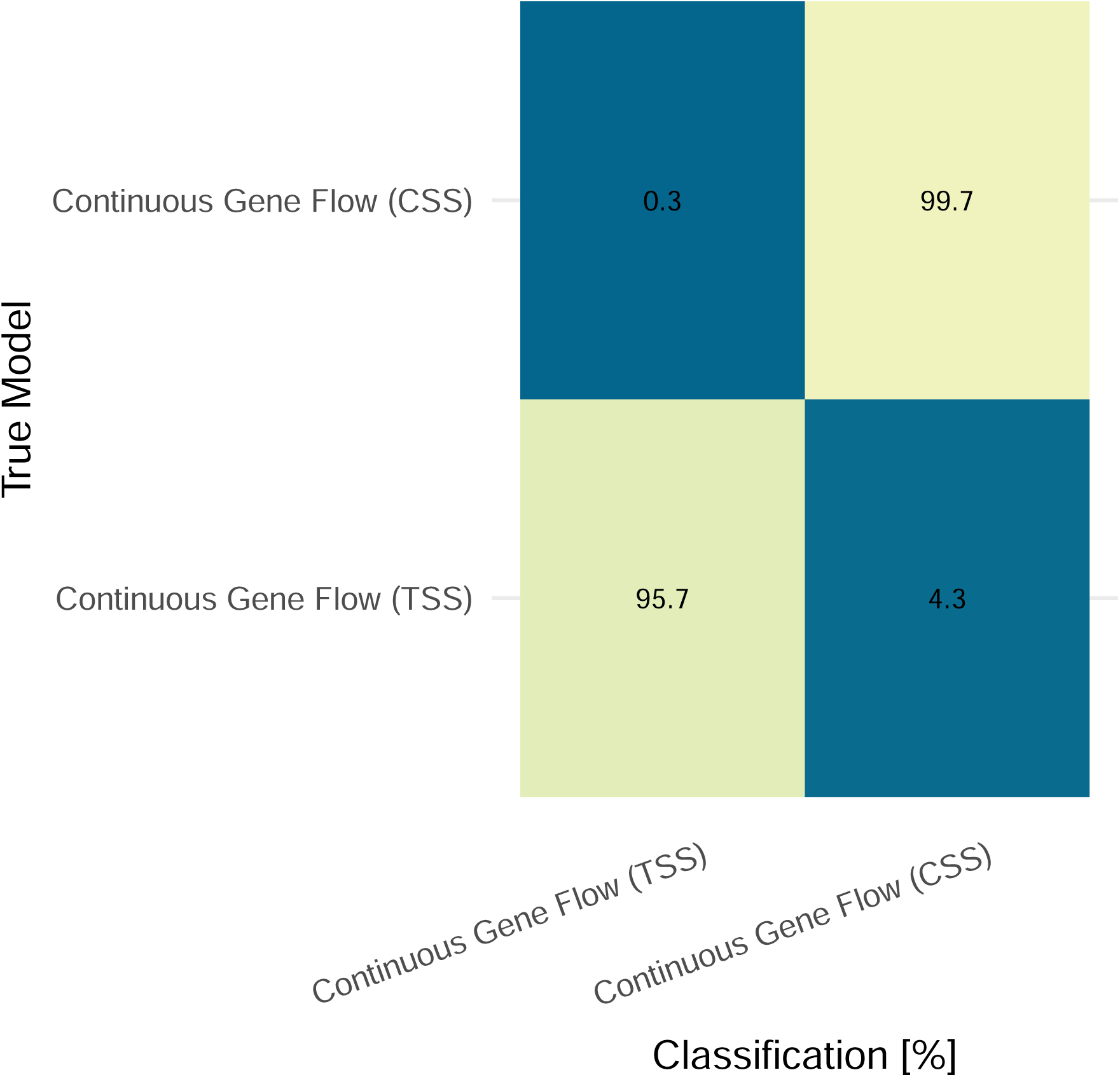
Confusion matrix for two continuous gene flow models under constant selfing (CSS) and transition-to-selfing scenario (TSS). For both models, simulated (out-of-bag) samples were classified by the trained random forest. Correct classifications are on the diagonal. Results are represented as percentages.

**Figure 14:**
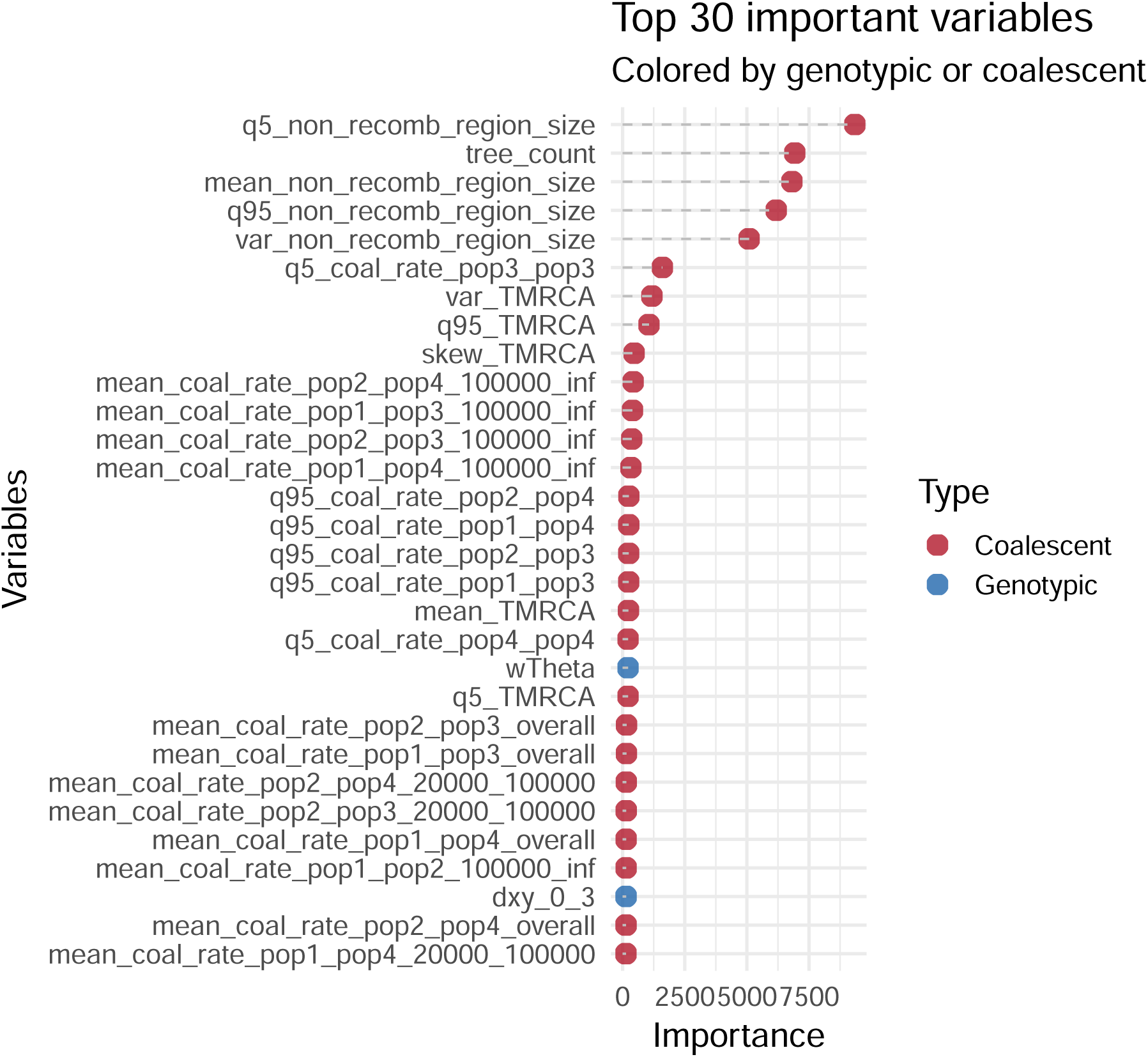
Variable importance for model classification. This plot highlights the top 30 variables that contributed most to the classification accuracy in the Approximate Bayesian Computation Random Forest (ABC-RF) framework for the continuous model in both, no-transition and transition set-up. Variables are ranked by their importance scores, with genotypic statistics (e.g., nucleotide diversity, fixation indices, site frequency spectrum metrics) distinguished from coalescent statistics (e.g., TMRCA moments, coalescent rates across time intervals, and tree counts). The dashed lines emphasize the importance gradient for each variable, with colors distinguishing variable types.

To visualize the footprint of transition to selfing in polymorphism data (Strütt et al., 2023), we ordered the 2,600 parameter sets by genomic position and highlighted the most important statistics in the ABC classifier. Regions classified as transition-to-selfing tended to have short non-recombining segments, high tree counts, and high TMRCA (Fig. 15). When one of these statistics is not extreme, the region was more likely classified under constant selfing. Most regions showed patterns consistent with constant selfing, suggesting that long segments of low-TMRCA trees might be dominant. This also implied that our simple transition model might not fully explain the real data.

**Figure 15:**
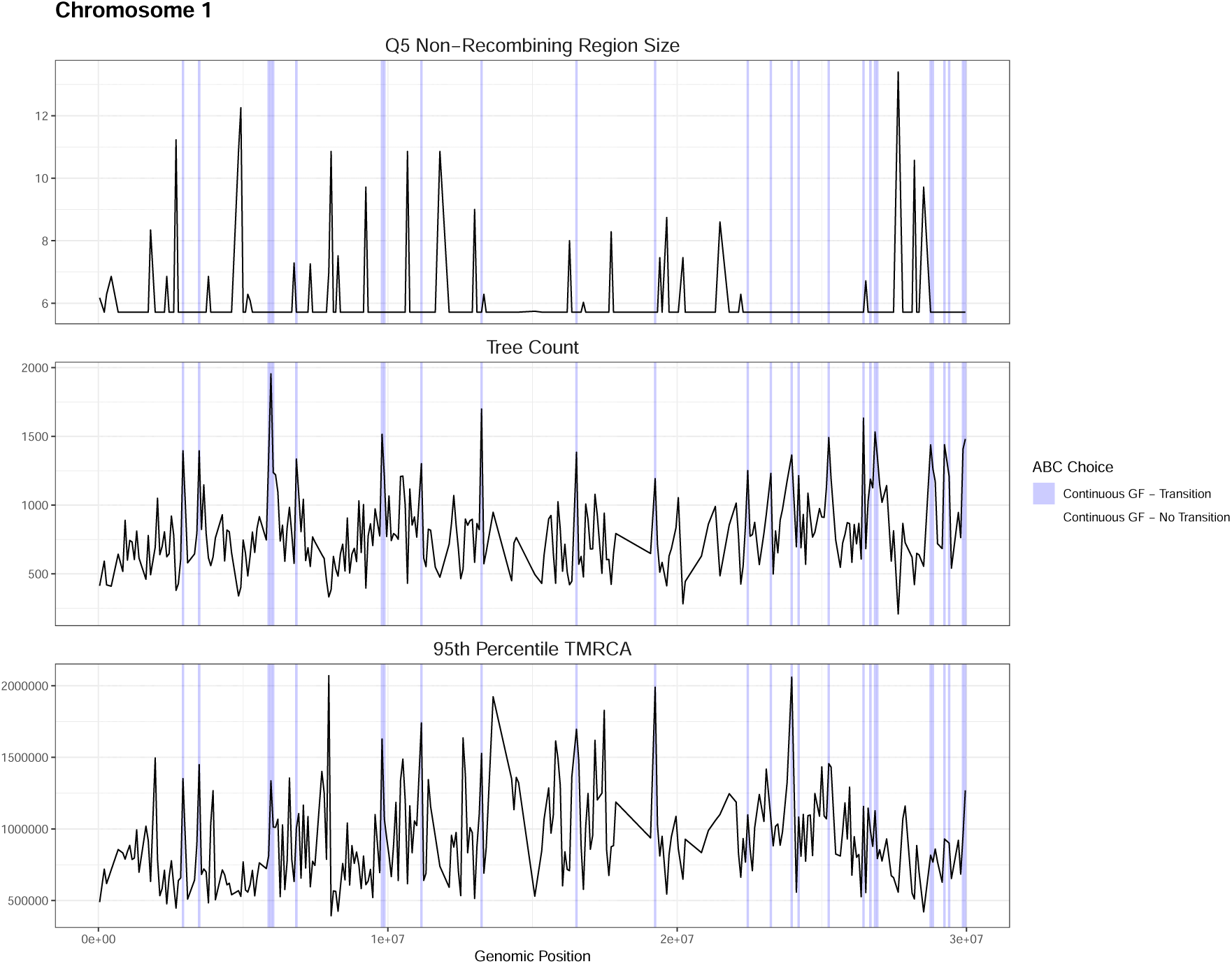
Genomic summary statistics and model choices across chromosomes for 2 demographic models. The figure displays genomic summary statistics across chromosome 1, highlighting regions associated with different demographic models. Lines represent the variation in tree count, mean non-recombining region size and 5 % quantile of non-recombining region size along genomic positions. Shaded areas indicate model choices derived from the ABC framework, with fill colors corresponding to specific demographic scenarios: Continuous gene flow with (in blue) and without transition scenario (in white). Each statistic is shown in a separate panel, scaled independently to facilitate comparison.

Repeating this procedure with six demographic models showed that secondary contact with transition and ancestral gene flow with transition were never chosen for any genomic region, so Fig. 16 focuses on four models: ancestral gene flow (no transition), continuous gene flow (with transition and without), and secondary contact (no transition). In these plots, the continuous gene flow model under transition again targets regions with extreme values for tree count, mean non-recombining segment size, and the 5 % quantile of non-recombining size. Theta Watterson did not show a simple monotonic association with model choice. Regions assigned to secondary contact without transition tended to show less extreme values for the displayed statistics. Overall, this suggests that explicitly including a transition-to-selfing scenario allows the ABC framework to capture part of the genomic heterogeneity in TMRCA and non-recombining segment structure that is not captured as well when only gene-flow models without a transition are considered.

**Figure 16:**
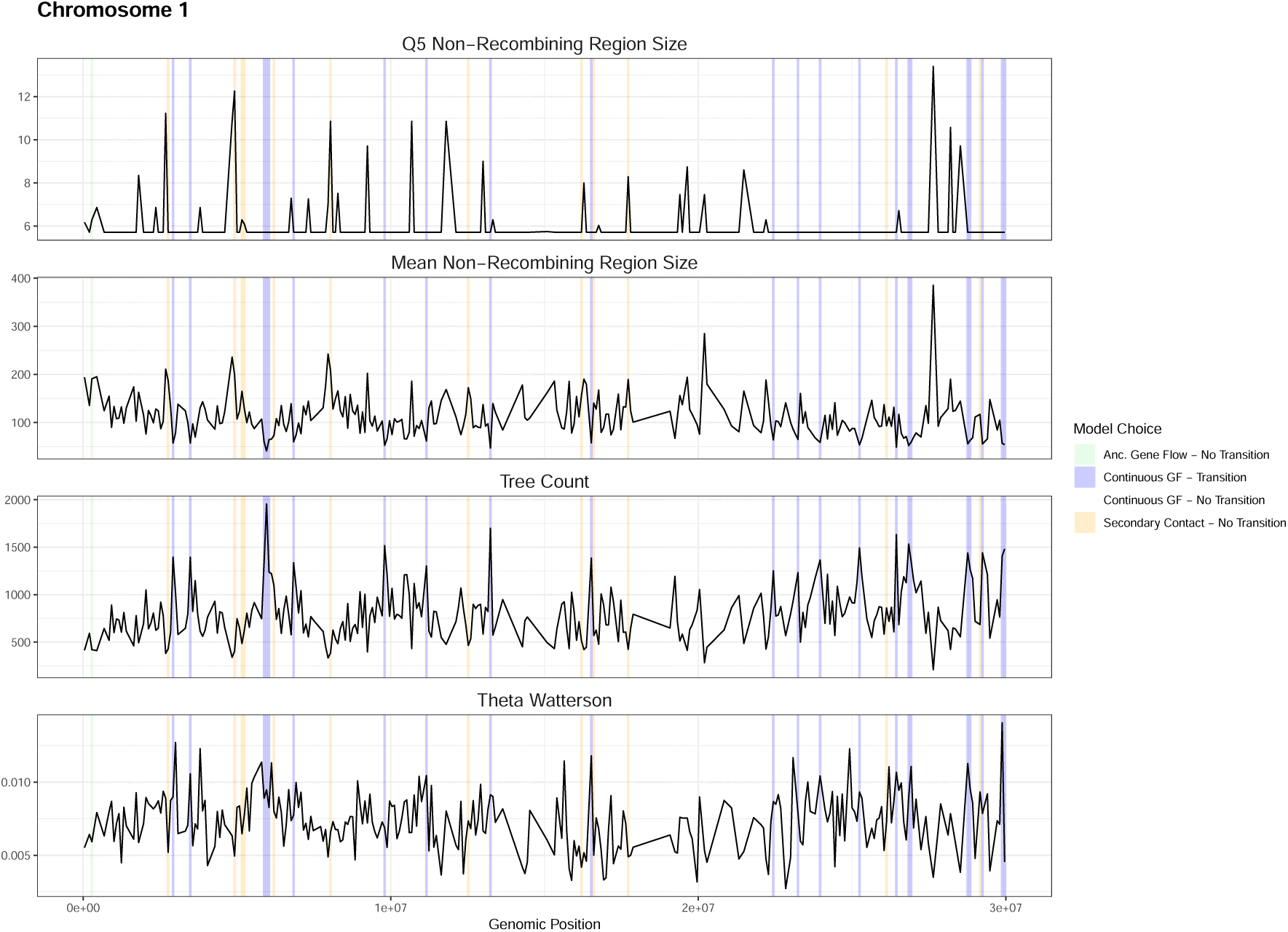
Genomic summary statistics and model choices across chromosomes for 4 demographic models. The figure displays genomic summary statistics across chromosome 1, highlighting regions associated with different demographic models. Lines represent the variation in tree count, mean non-recombining region size, 5 % quantile of non-recombining region size, and Theta Watterson along genomic positions. Shaded areas indicate model choices derived from the ABC framework, with fill colors corresponding to specific demographic scenarios: ancestral gene flow without transition scenario (in green), continuous gene flow with (in blue) and without transition scenario (in white), and secondary contact without transition (in yellow). Each statistic is shown in a separate panel, scaled independently to facilitate comparison.

## 4 Discussion

We show that ABC models of demographic history that include introgression can yield misleading results when they fail to account for a recent transition to self-fertilization. Particularly, when a constant selfing rate was assumed, the ABC model misclassified models more frequently when the true history included a selfing transition, particularly under scenarios of continuous gene flow or secondary contact. Conversely, assuming a recent shift to selfing when none occurred introduced errors under models with no gene flow or ancient gene flow. As many plants have transitioned to selfing in their evolutionary history (Mattila et al., 2020) and may have also experienced introgression, demographic events could be misinterpreted. Similar challenges arise from factors such as population structure (Durand et al., 2011; Mazet et al., 2016; Tournebize & Chikhi, 2024), natural selection (Johri et al., 2022; Smith & Hahn, 2024), or large divergence with heterogeneous rates (Koppetsch et al., 2024; Xiong et al., 2022). Accurately inferring introgression histories is crucial for understanding modes of speciation. Numerous studies supporting gene flow between closely related populations interpret this as speciation with subsequent gene flow (Monnet et al., 2025), especially in selfing plant species (Brandvain et al., 2014; Dittberner et al., 2022; Rahnamae et al., 2026). The results of this study highlight the need for caution when analyzing gene flow history (Monnet et al., 2025), particularly when a recent transition to selfing is involved. However, it is still unknown how common such biases are likely to be in natural systems, because a systematic/exhaustive study of the occurrence of changes in life-history traits has not yet been performed (Metzger et al., 2026).

Importantly, we did not simulate other factors that could bias inference, such as background selection or varying recombination and mutation rates. Background selection is expected to be particularly significant in selfing species due to increased genome-wide homozygosity. This results in a higher proportion of recombination events occurring between nearly identical chromosomes, effectively reducing the recombination rate (Nordborg, 2000). Consequently, background selection (Charlesworth et al., 1993; Dickinson et al., 2003) and selective sweeps (Hartfield & Bataillon, 2020) are more pronounced. Neglecting background selection can lead to underestimations of population size and the misinterpretation of recent population growth. Coalescent trees in regions affected by background selection are typically shorter (as in small populations) and have longer terminal branches (as in expanding populations). To understand its impact, future simulations should incorporate background selection to evaluate its effect on model choice and error rates. However, Strütt et al. (2023) have shown that background selection does not significantly affect the estimation of a transition to selfing. Therefore, we assume that the estimates remain accurate in both the ABC and teSMC approaches. For real data, as shown in Rahnamae et al., 2026, there is no clear overlap between introgressions and selective sweeps, suggesting minimal influence of positive selection on model choice for *A. sagittata* and *A. nemorensis*.

We used teSMC to date the transition to selfing in *A. sagittata* and *A. nemorensis* and identified an event occurring 470,000–890,000 years ago (depending on chromosome). In comparison, the ABC parameter estimation detected a transition to selfing at approximately 1,464,935 years ago. The difference likely arises because the ABC estimation was based on a single region characterized as an outcrossing region with TMRCAs consistently older than the expected time of the transition to selfing. As a result, it provides an older baseline due to its predominantly ancient TMRCAs. The ABC estimate also exceeds the calculated divergence time of the *Arabis hirsuta* clade (approximately 1.2 million years ago; Karl and Koch, 2014), which includes both *A. sagittata* and *A. nemorensis*. Therefore, we assume that the teSMC estimate is likely closer to the true timing of the transition, though further analysis should include additional genomic regions to refine it. Moreover, the teSMC timeframe overlaps with the transition to selfing in *Arabidopsis thaliana* (approximately 590,000 to 750,000 years ago; Strütt et al., 2023) and the Middle Pleistocene Transition (approximately 1,250,000 to 650,000 years ago), a period of extended and extreme glacial cycles (Tzedakis et al., 2017; Willeit et al., 2019). These environmental changes were linked to increased speciation events (Willi et al., 2022), and we speculate that self-fertilization may have evolved also as an adaptive strategy for several species of the *Arabis* clade. Our study demonstrates the usefulness of full-genome polymorphism data for evaluating the potential influence of environmental changes on the evolution of life-history traits (Metzger et al., 2026).

Our study is among the first to utilize tree sequences within an ABC framework, leveraging underlying genealogies for downstream inferences. While tree sequences have gained popularity in simulations (Baumdicker et al., 2021; Haller & Messer, 2023; Kelleher et al., 2016, 2018), population genetic inference has primarily relied on low-dimensional summaries of ancestral recombination graphs (ARGs), such as site frequency spectra or F_ST_, rather than the full ARG (Korfmann et al., 2023, 2026). Advances now allow ARGs to be directly recovered for demographic inference (Deng et al., 2024; Korfmann et al., 2026). Using tree sequences enables improved downstream inference of recent demographic events, such as natural selection, recent admixture events and introgression (Speidel et al., 2025; Whitehouse et al., 2024). However, inferred ARGs may contain errors, and these errors are not accounted for when applying ABC to observed datasets. Ideally, methods like SINGER (Deng et al., 2024) or cxt (Korfmann et al., 2026) could be applied to simulated data, incorporating errors directly into the training of the ABC classifier. This approach involves inferred trees for the simulated data, rather than using the true trees directly. While we only tested a subset of simulations comparing true and inferred tree sequences, these comparisons highlight potential biases in the observed data for *A. sagittata* and *A. nemorensis*.

From Figures 15 and 16, comparing the transition-to-selfing scenario (TSS) with the constant selfing scenario (CSS) identifies characteristic genomic regions which ancestry goes back to the outcrossing phase. Previously, transition matrices alone were assumed to achieve this (Strütt et al., 2023). Figure 14 shows that coalescent statistics are the primary contributors for detecting transitions. However, our ABC framework treats genomic windows independently, selecting the model with the most window votes. This limitation favors CSS for *A. sagittata* and *A. nemorensis* since regions with selfing signatures (e.g., long stretches with no recombination and more recent TMRCA) dominate over smaller transition (outcrossing) regions. Future versions of the ABC framework should incorporate transition matrix features (Strütt et al., 2023) and adjust for varying transition probabilities across genomic regions. An alternative approach would estimate transition timing directly from TMRCA distributions and compare regions classified as transitions with those without transitions in simulated data (extending the work of Korfmann et al. (2026)). Additionally, constructing phylogenies for S-locus regions in both species could complement these efforts, although biases may persist if only a small subset of genetic variants is analyzed (Strütt et al., 2023), and also this can be difficult if the S-locus region is absent.

Dittberner et al. (2022) reported slightly lower migration rates compared to our findings. This deviation may arise from two factors: our analysis currently focuses on a single genomic region, and we have not yet included asymmetric gene flow, potentially leading to the overestimation of migration rates. Additionally, Dittberner et al. (2022) identified secondary contact as the best-fitting demographic history, characterized by ancient introgression together with contemporary gene flow. In contrast, our results favor a continuous gene flow model, as a result the parameter estimation is not based on the same demographic model and may be therefore different. The discrepancy between demographic models can be explained by several factors. First, by accounting for a transition to selfing, our approach better distinguishes between secondary contact and continuous gene flow models, as shown by simulations in Fig. 4. Second, we use here coalescent statistics. Notably, when using only genotypic statistics and the constant selfing scenario (CSS), our results favor the continuous gene flow model. However, the secondary contact model also receives significant support (45 % versus 33 %), suggesting that genotypic statistics alone struggle to fully differentiate between these models. Differences in simulation settings further contribute to the discrepancies. Dittberner et al. (2022) conducted 50,000 simulations, each comprising 500 regions of 75 kb, while we performed 10,000 simulations with regions spanning 2 Mb. Additionally, our analysis using genotypic statistics relied on a subset of 65 summary statistics, whereas Dittberner et al. (2022) used 236 statistics. Despite these differences, our simulations confirm that inference problems can arise when relying on incorrect mating system models. By accounting for the transition to selfing, we are confident in identifying the continuous gene flow model as the correct demographic history.

Moving forward, we recommend interpreting demographic inferences about gene flow with caution when a transition to selfing may have occurred, particularly if it is recent. The results presented here are derived from a small set of gene flow models and selfing histories, and thus, are insufficient to adequately cover all possible effects on demographic inferences. Models that ignore a transition to selfing may influence interpretations about the presence, direction, and timing of introgression or speciation with gene flow. We develop an ABC framework that incorporates genotypic and coalescent summary statistics, which improves inference accuracy. Our findings are consistent with Korfmann et al. (2026) and Whitehouse et al. (2024), showing similar improvements in using deep learning methods, and validate the power of this approach within an ABC framework. The inclusion of tree sequences for downstream inference leads to the detection of genomic signals that genotypic statistics alone might have missed. We encourage further evaluation of tree sequence-based methods as a flexible and powerful tool for improving ABC inferences.

## Acknowledgements - Financial support

The authors acknowledge support by the Deutsche Forschungsgemeinschaft - German Research Council (DFG) project 447121532. The funder had no role in study design, data collection and analysis, decision to publish, or preparation of the manuscript.

## Data availability statement

There is no data associated with this manuscript.

## Supplementary figures

**Figure 17:**
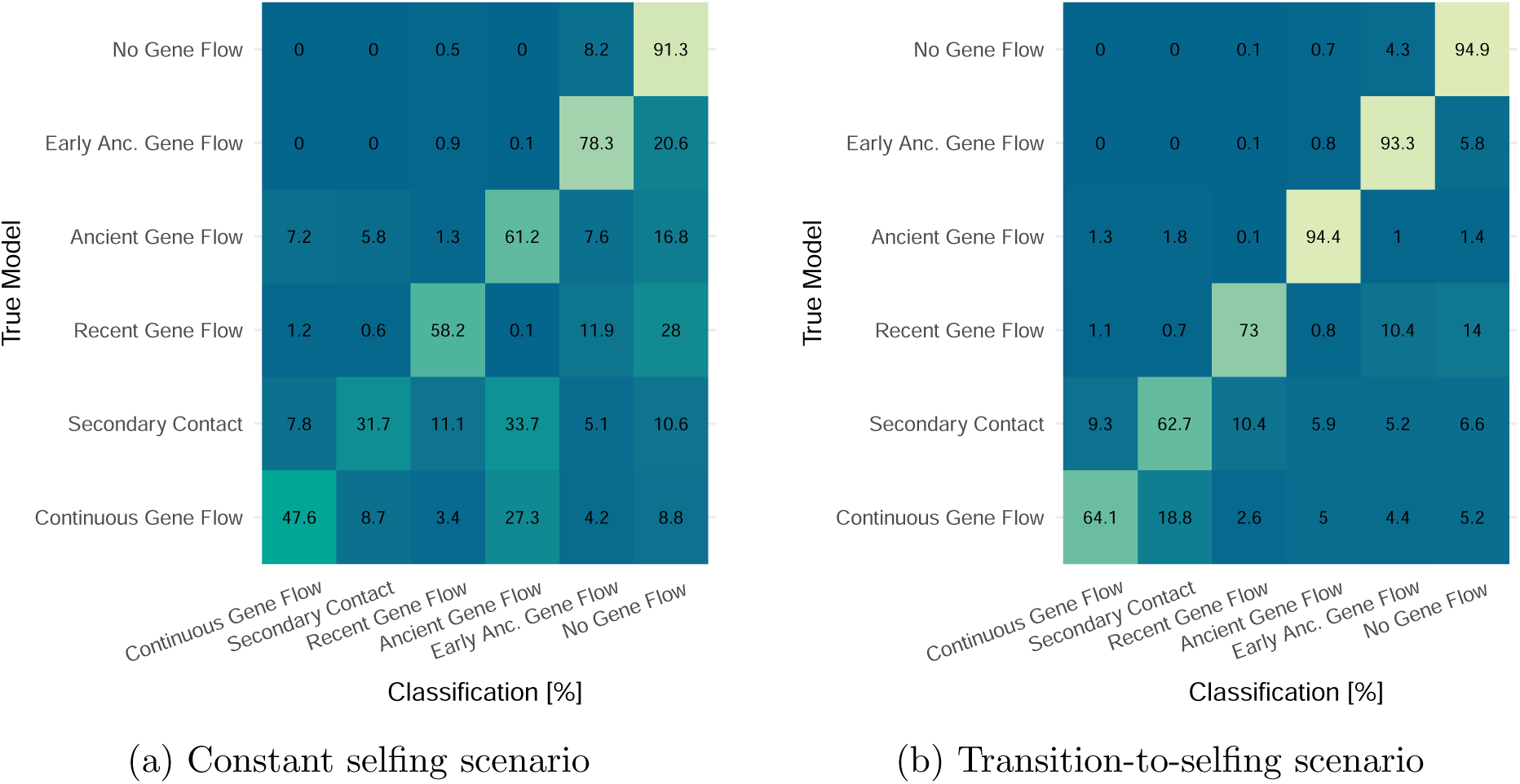
Confusion matrix for model classification using ABC-RF with 200,000 bp windows. The confusion matrix summarizes the classification accuracy of the ABC-RF model, comparing the predicted models (x-axis) against the true models (y-axis). Values represent the percentage of correct and misclassified instances for each model, calculated as row-wise proportions. The diagonal cells indicate the percentage of correctly classified instances for each true model, while off-diagonal cells represent misclassifications. Colors reflect classification percentages, with darker shades indicating higher accuracy. This matrix is derived from out-of-bag (OOB) predictions, which validate the model’s performance by using samples excluded from the bootstrap resampling process during model training, ensuring unbiased accuracy estimates.

**Figure 18:**
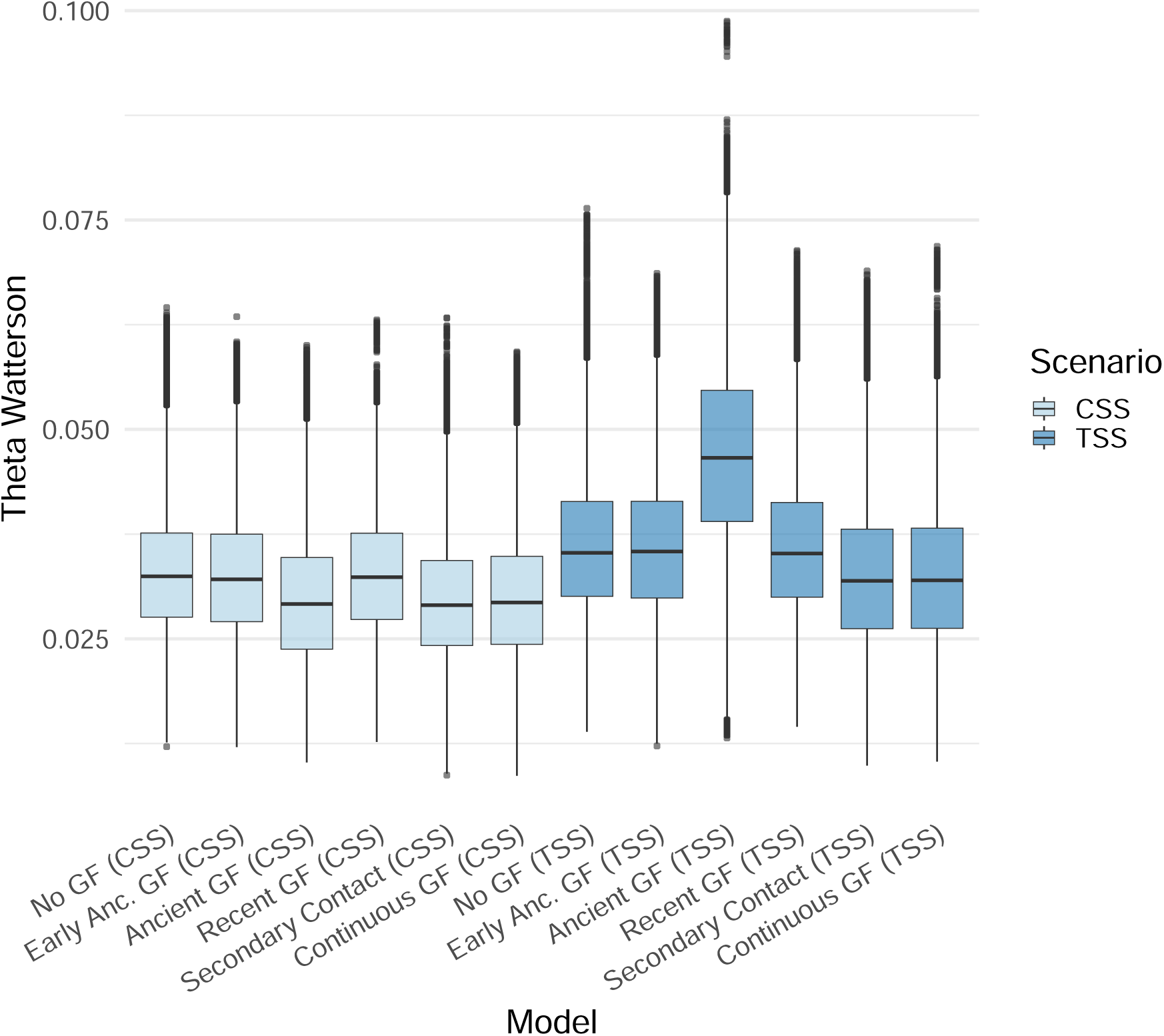
Boxplot showing the distribution of Watterson’s Theta (Θ*_W_*) across different demographic models. The models are grouped by scenario (CSS or TSS) and further categorized into specific demographic models. Colors represent the scenario types (CSS: Constant selfing scenarios, TSS: Transition-to-selfing scenario). The x-axis represents the demographic models, and the y-axis shows Watterson’s Theta values.

**Figure 19:**
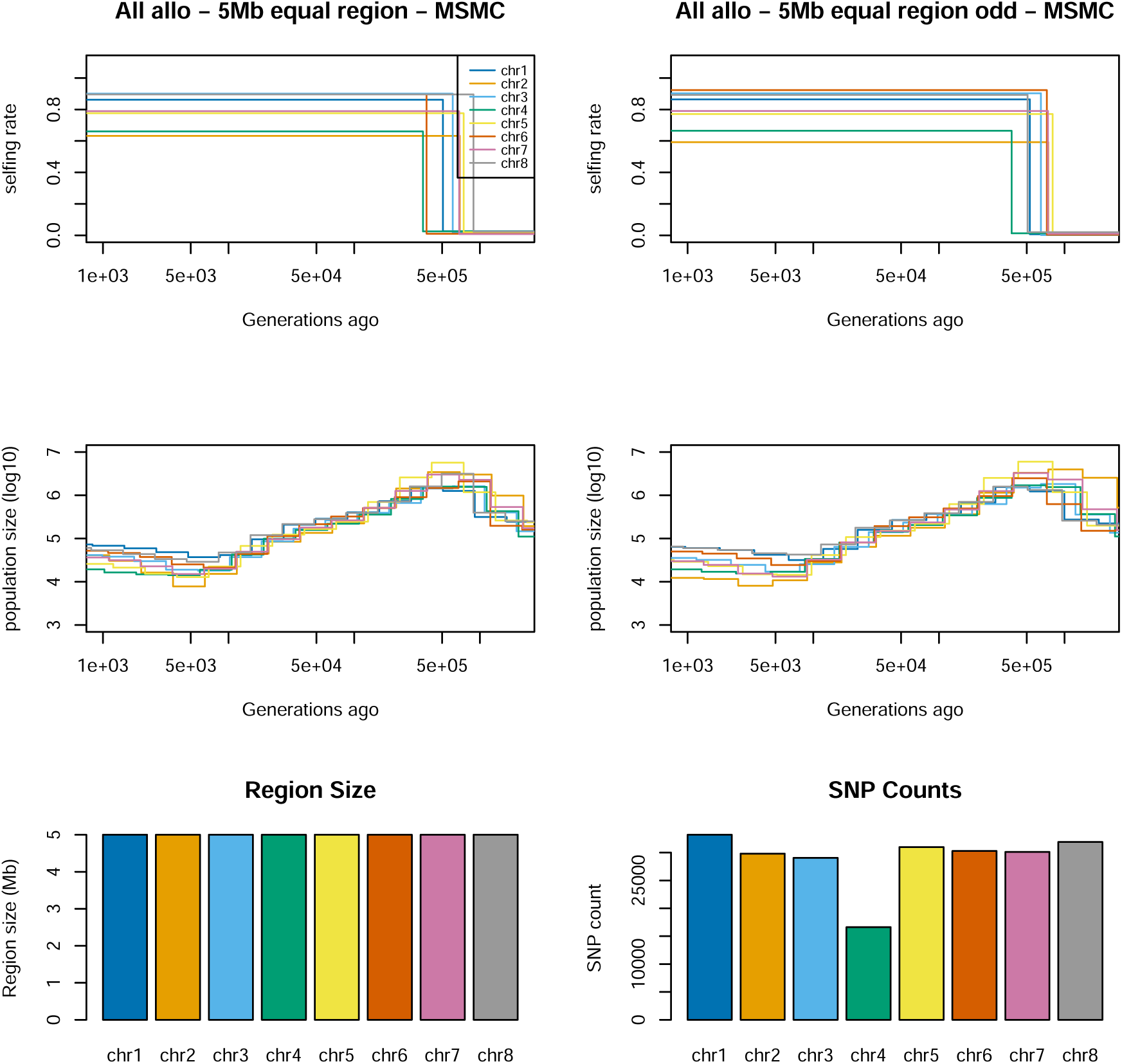
Demographic inference and the transition from outcrossing to selfing in allopatric *A. nemorensis* and *A. sagittata* for 5 Mb regions. The top row depicts transition times and selfing rates inferred for 5 Mb regions across all chromosomes in both species. The middle row presents estimated population sizes over time using teSMC, with population size estimates derived from MSMC2 time intervals. The bottom row shows the size of the genomic regions alongside their respective SNP counts for each chromosome. The analysis was conducted using 12 pseudo-diploid individuals, each combining two haploid sequences from distinct individuals of the same species, and included 12 haploid samples from both *A. nemorensis* and *A. sagittata*. The two columns in the first two rows represent different sets of pseudo-diploid individuals. Colored lines in each plot represent individual chromosomes.

**Figure 20:**
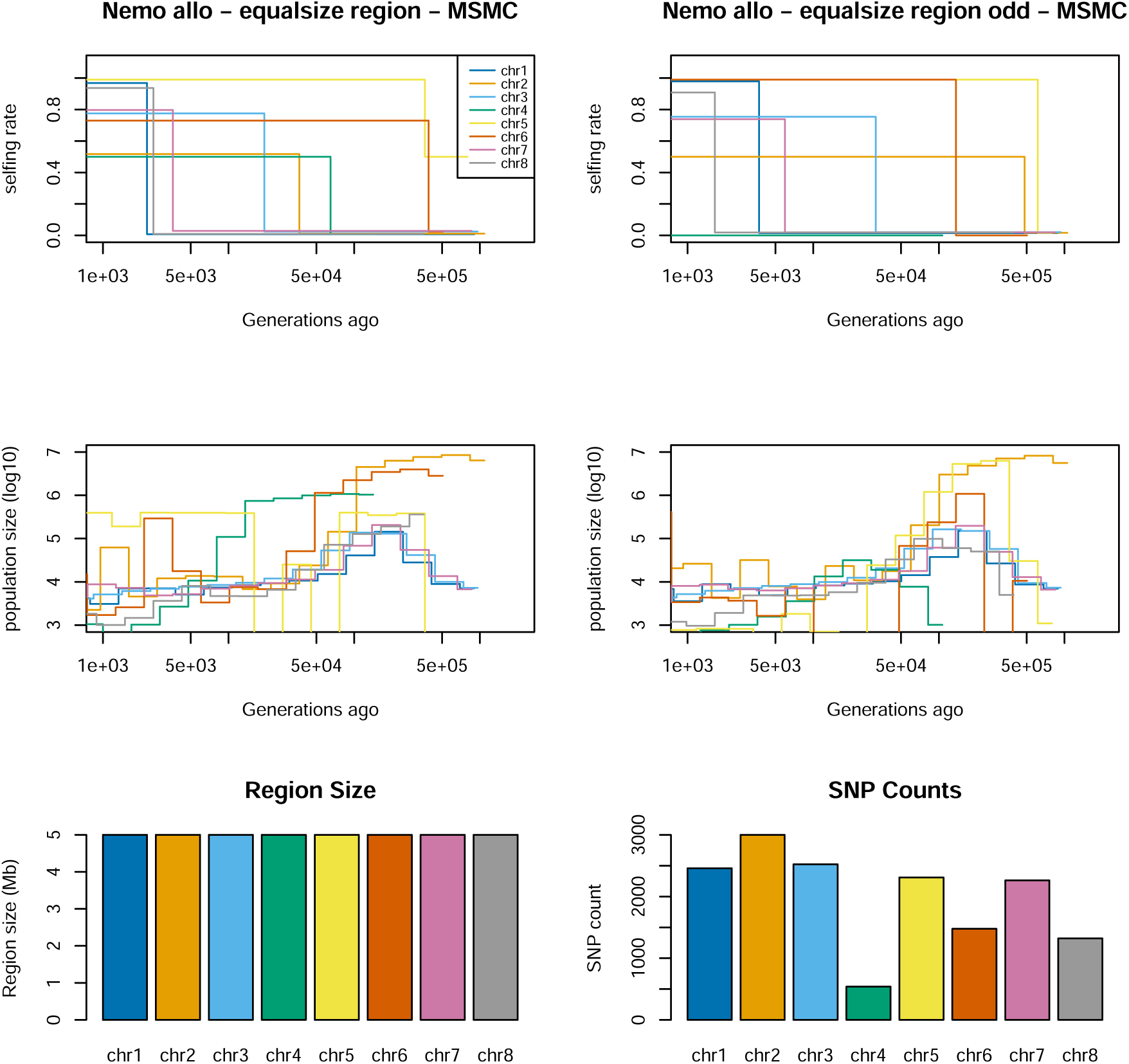
Demographic inference and the transition from outcrossing to selfing in allopatric *A. nemorensis* for 5 Mb regions. The top row depicts transition times and selfing rates inferred for 5 Mb regions across all chromosomes in *A. nemorensis*. The middle row presents estimated population sizes over time using teSMC, with population size estimates derived from MSMC2 time intervals. The bottom row shows the size of the genomic regions alongside their respective SNP counts for each chromosome. The analysis was conducted using 6 pseudo-diploid individuals, each combining two haploid sequences from distinct individuals of the same species, and included 6 haploid samples from *A. nemorensis*. The two columns in the first two rows represent different sets of pseudo-diploid individuals. Colored lines in each plot represent individual chromosomes.

**Figure 21:**
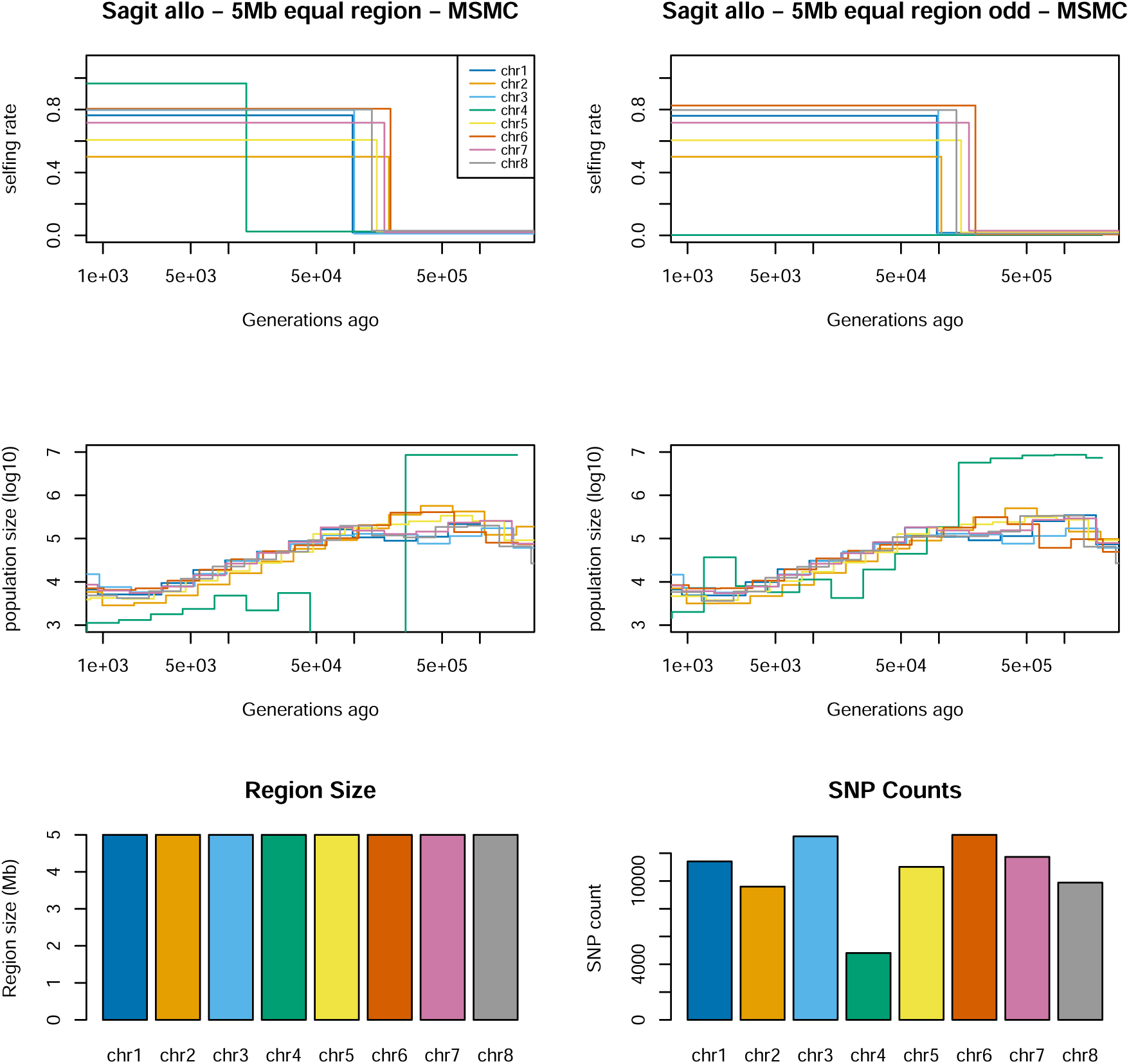
Demographic inference and the transition from outcrossing to selfing in allopatric *A. sagittata* for 5 Mb regions. The top row depicts transition times and selfing rates inferred for 5 Mb regions across all chromosomes in *A. sagittata*. The middle row presents estimated population sizes over time using teSMC, with population size estimates derived from MSMC2 time intervals. The bottom row shows the size of the genomic regions alongside their respective SNP counts for each chromosome. The analysis was conducted using 6 pseudo-diploid individuals, each combining two haploid sequences from distinct individuals of the same species, and included 6 haploid samples from *A. nemorensis*. The two columns in the first two rows represent different sets of pseudo-diploid individuals. Colored lines in each plot represent individual chromosomes.

**Figure 22:**
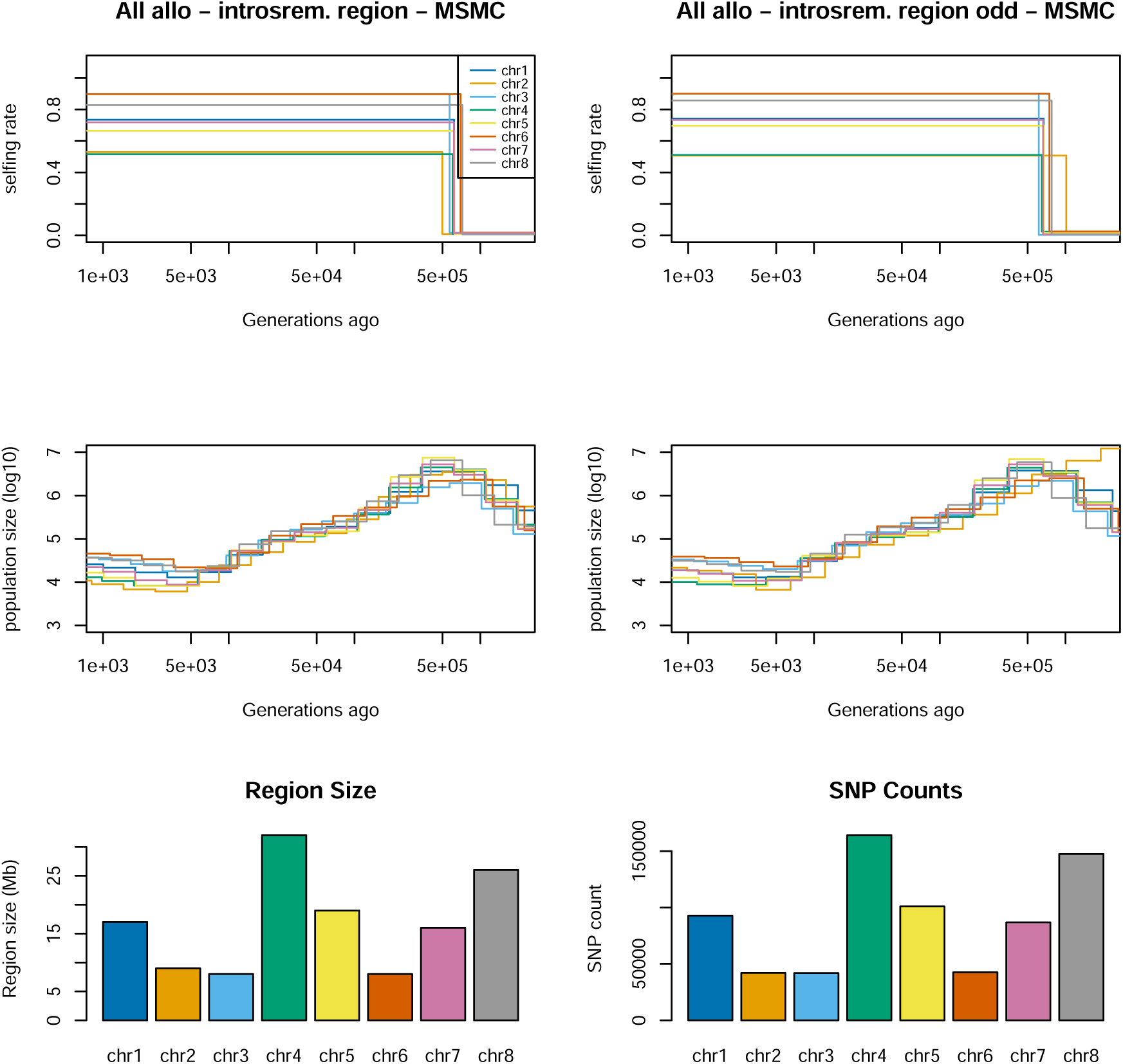
Demographic inference and the transition from outcrossing to selfing in allopatric *A. nemorensis* and *A. sagittata* for introsrem regions. The top row depicts transition times and selfing rates inferred for another genomic region across all chromosomes in both species. These regions vary in size between chromosomes. The middle row presents estimated population sizes over time using teSMC, with population size estimates derived from MSMC2 time intervals. The bottom row shows the size of the genomic regions alongside their respective SNP counts for each chromosome. The analysis was conducted using 12 pseudo-diploid individuals, each combining two haploid sequences from distinct individuals of the same species, and included 12 haploid samples from both *A. nemorensis* and *A. sagittata*. The two columns in the first two rows represent different sets of pseudo-diploid individuals. Colored lines in each plot represent individual chromosomes.

**Figure 23:**
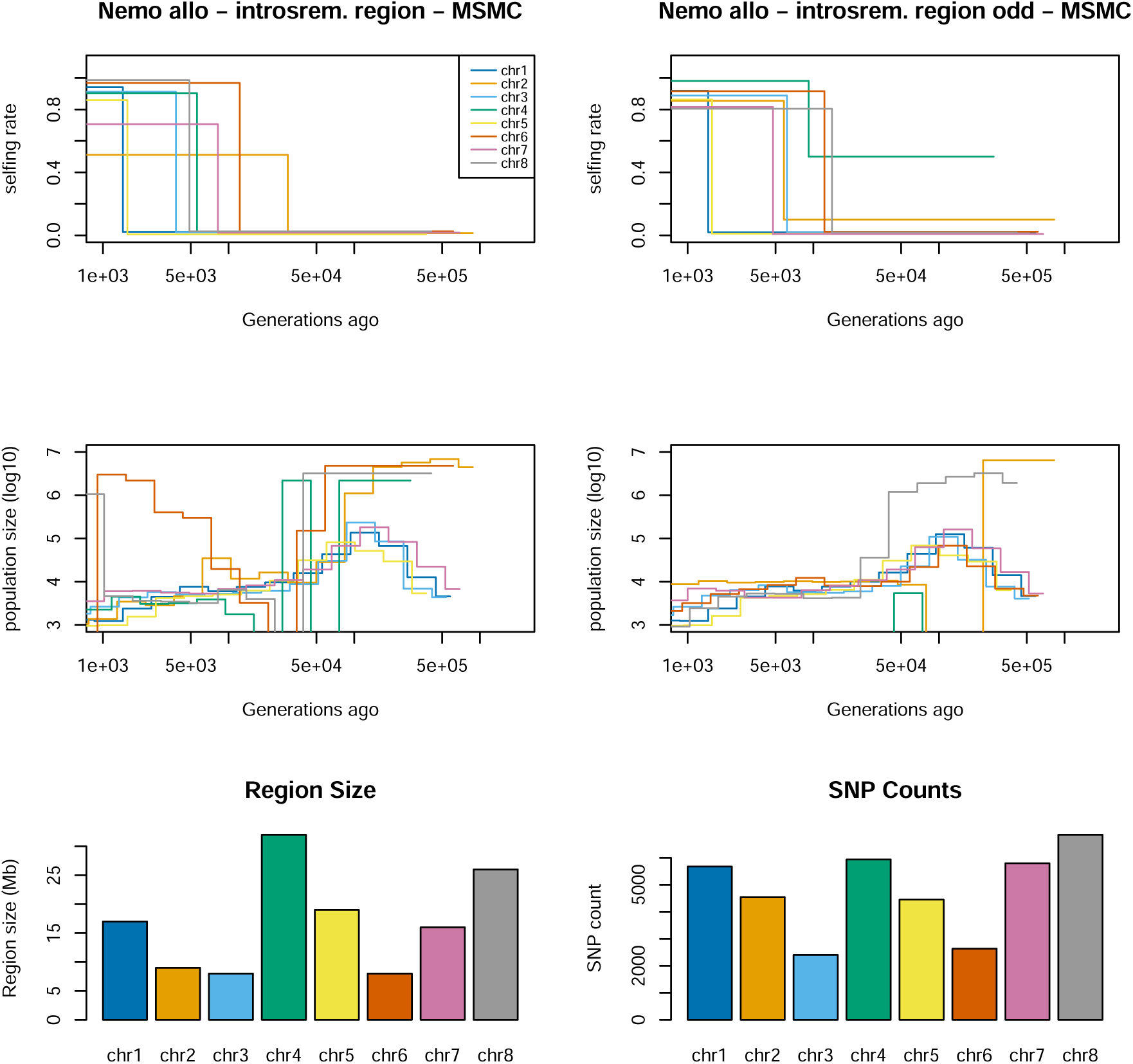
Demographic inference and the transition from outcrossing to selfing in allopatric *A. nemorensis* for introsrem regions. The top row depicts transition times and selfing rates inferred for another genomic region across all chromosomes in both species. These regions vary in size between chromosomes. The middle row presents estimated population sizes over time using teSMC, with population size estimates derived from MSMC2 time intervals. The bottom row shows the size of the genomic regions alongside their respective SNP counts for each chromosome. The analysis was conducted using 6 pseudo-diploid individuals, each combining two haploid sequences from distinct individuals of the same species, and included 6 haploid samples from both *A. nemorensis*. The two columns in the first two rows represent different sets of pseudo-diploid individuals. Colored lines in each plot represent individual chromosomes.

**Figure 24:**
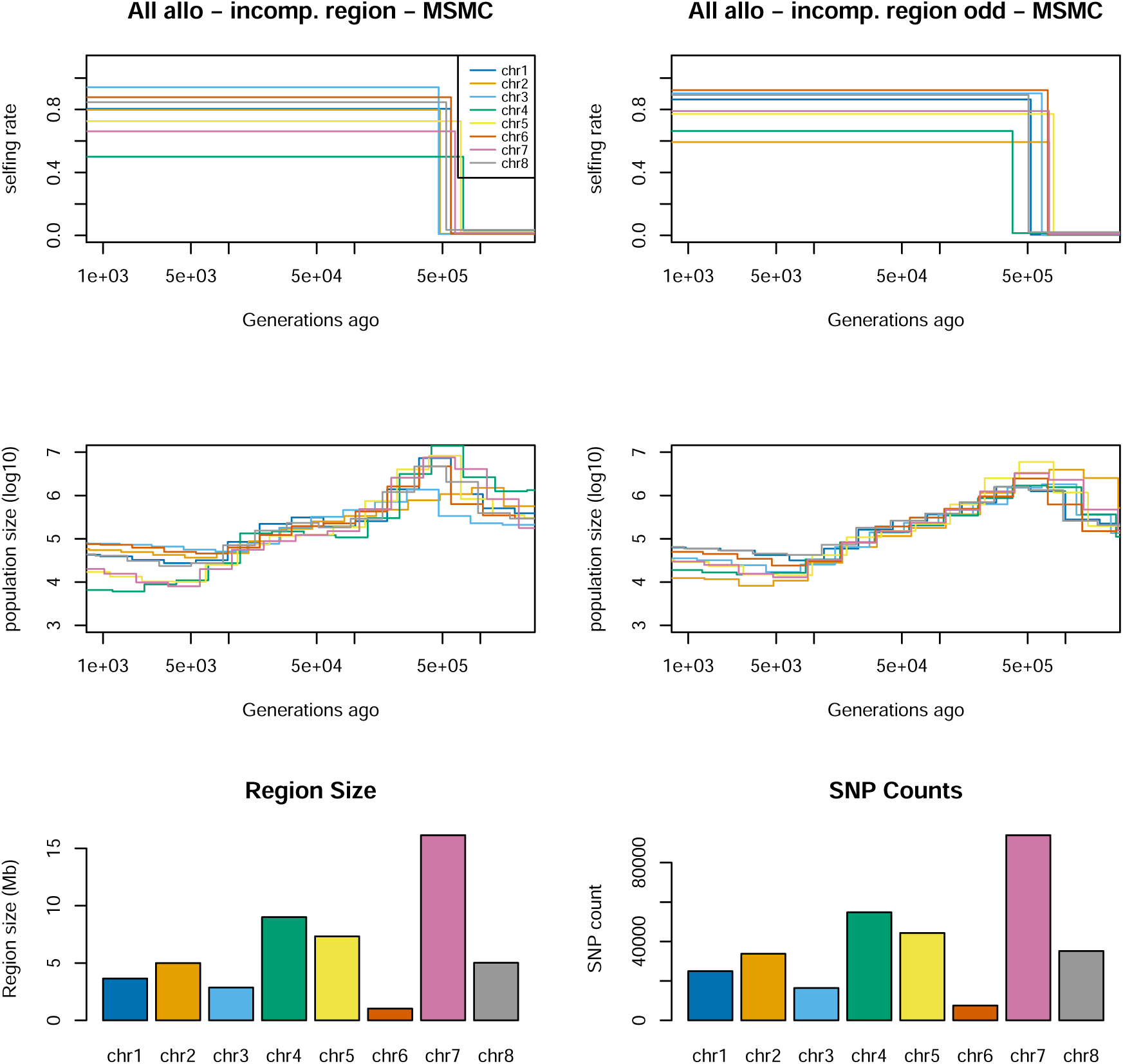
Demographic inference and the transition from outcrossing to selfing in allopatric *A. nemorensis* and *A. sagittata* for incomp regions. The top row depicts transition times and selfing rates inferred for another genomic region across all chromosomes in both species. These regions vary in size between chromosomes. The middle row presents estimated population sizes over time using teSMC, with population size estimates derived from MSMC2 time intervals. The bottom row shows the size of the genomic regions alongside their respective SNP counts for each chromosome. The analysis was conducted using 12 pseudo-diploid individuals, each combining two haploid sequences from distinct individuals of the same species, and included 12 haploid samples from both *A. nemorensis* and *A. sagittata*. The two columns in the first two rows represent different sets of pseudo-diploid individuals. Colored lines in each plot represent individual chromosomes.

**Figure 25:**
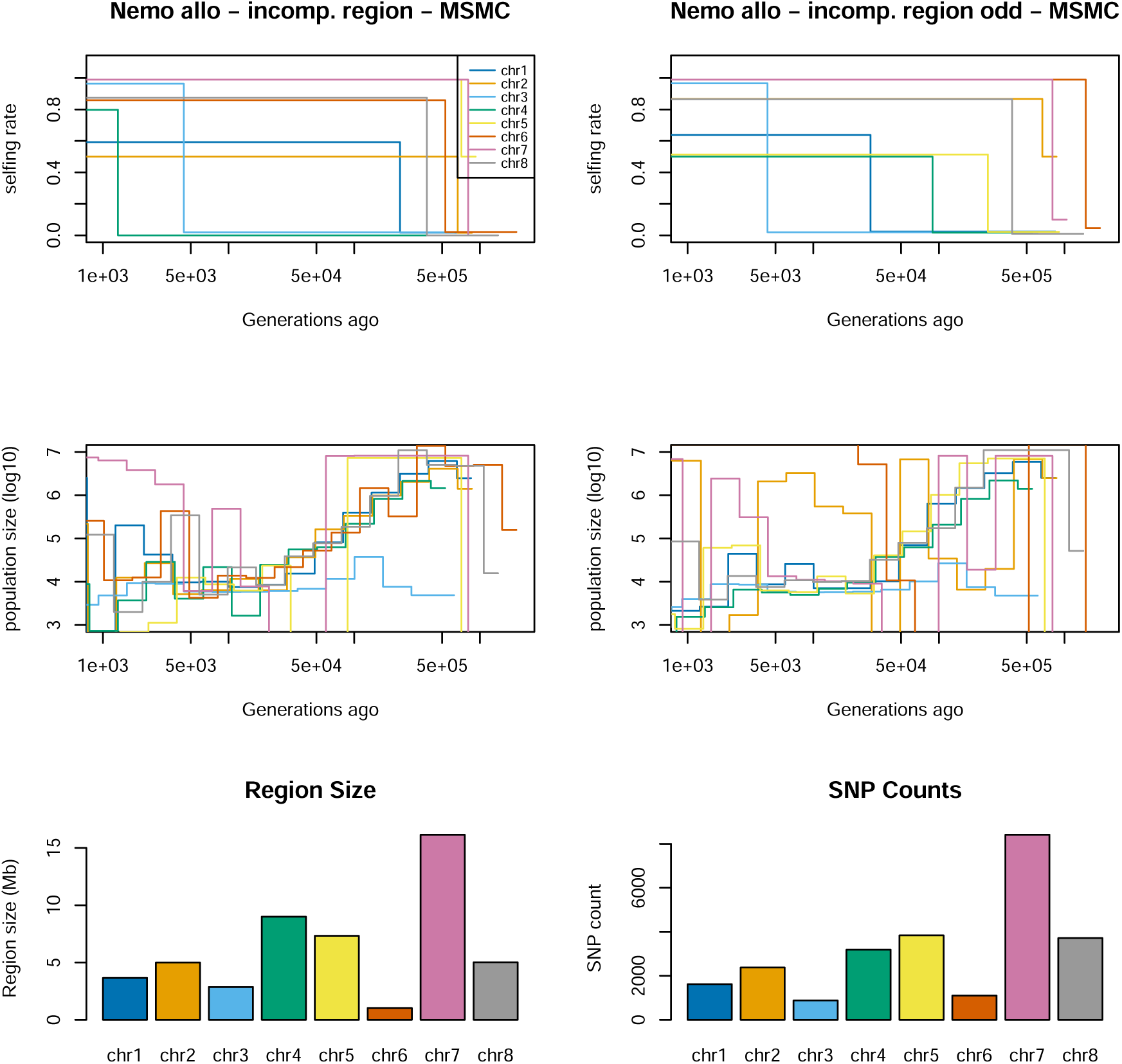
Demographic inference and the transition from outcrossing to selfing in allopatric *A. nemorensis* for incomp regions. The top row depicts transition times and selfing rates inferred for another genomic region across all chromosomes in both species. These regions vary in size between chromosomes. The middle row presents estimated population sizes over time using teSMC, with population size estimates derived from MSMC2 time intervals. The bottom row shows the size of the genomic regions alongside their respective SNP counts for each chromosome. The analysis was conducted using 6 pseudo-diploid individuals, each combining two haploid sequences from distinct individuals of the same species, and included 6 haploid samples from both *A. nemorensis*. The two columns in the first two rows represent different sets of pseudo-diploid individuals. Colored lines in each plot represent individual chromosomes.

**Figure 26:**
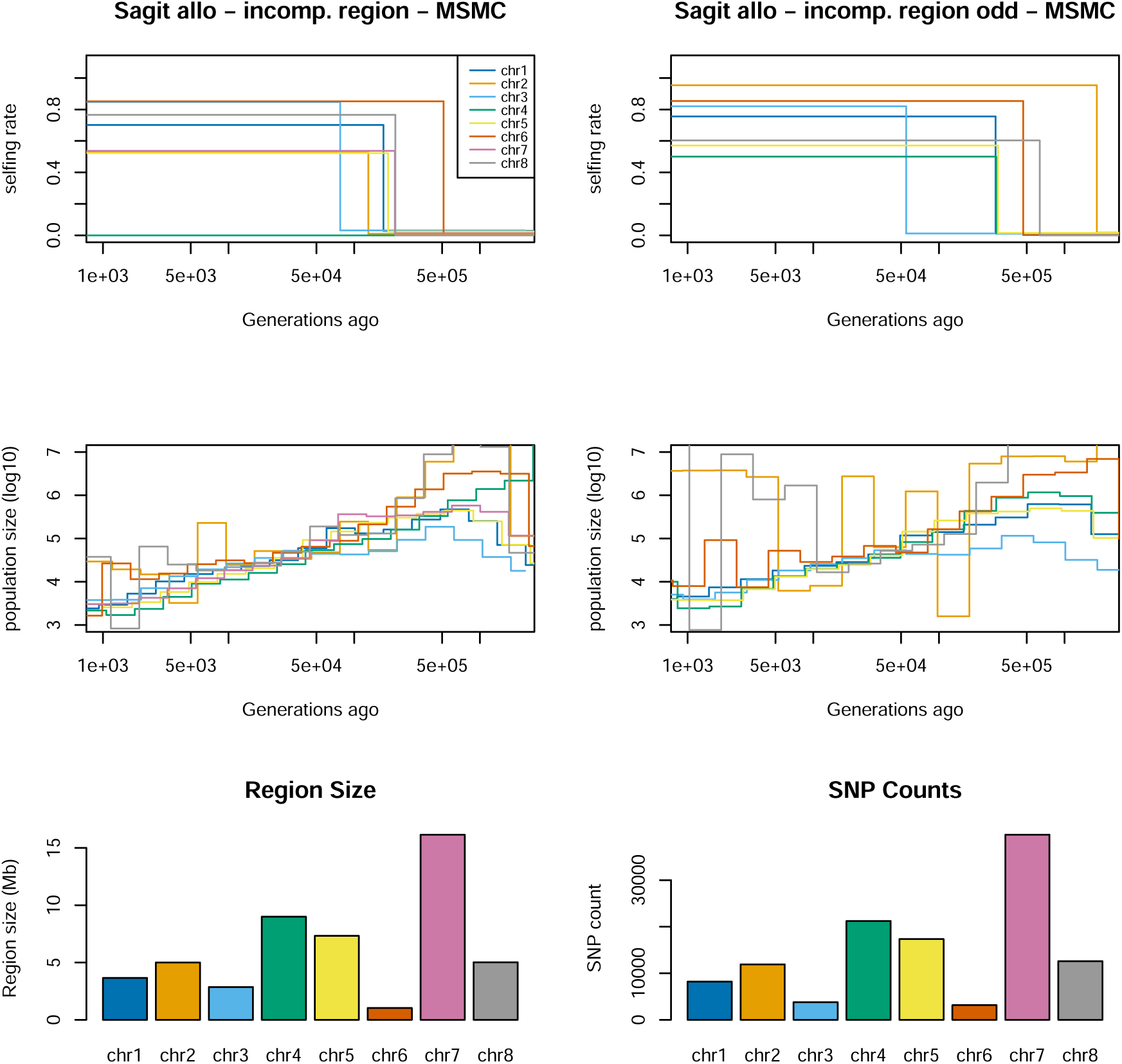
Demographic inference and the transition from outcrossing to selfing in allopatric *A. sagittata* for incomp regions. The top row depicts transition times and selfing rates inferred for another genomic region across all chromosomes in both species. These regions vary in size between chromosomes. The middle row presents estimated population sizes over time using teSMC, with population size estimates derived from MSMC2 time intervals. The bottom row shows the size of the genomic regions alongside their respective SNP counts for each chromosome. The analysis was conducted using 6 pseudo-diploid individuals, each combining two haploid sequences from distinct individuals of the same species, and included 6 haploid samples from both *A. sagittata*. The two columns in the first two rows represent different sets of pseudo-diploid individuals. Colored lines in each plot represent individual chromosomes.

